# RERconverge Expansion: Using Relative Evolutionary Rates to Study Complex Categorical Trait Evolution

**DOI:** 10.1101/2023.12.06.570425

**Authors:** Ruby Redlich, Amanda Kowalczyk, Michael Tene, Heather H. Sestili, Kathleen Foley, Elysia Saputra, Nathan Clark, Maria Chikina, Wynn K. Meyer, Andreas Pfenning

**Affiliations:** Carnegie Mellon University; Lehigh University; University of Pittsburgh

## Abstract

Comparative genomics approaches seek to associate evolutionary genetic changes with the evolution of phenotypes across a phylogeny. Many of these methods, including our evolutionary rates based method, RERconverge, lack the capability of analyzing non-ordinal, multicategorical traits. To address this limitation, we introduce an expansion to RERconverge that associates shifts in evolutionary rates with the convergent evolution of multi-categorical traits. The categorical RERconverge expansion includes methods for performing categorical ancestral state reconstruction, statistical tests for associating relative evolutionary rates with categorical variables, and a new method for performing phylogenetic permulations on multi-categorical traits. In addition to demonstrating our new method on a three-category diet phenotype, we compare its performance to naive pairwise binary RERconverge analyses and two existing methods for comparative genomic analyses of categorical traits: phylogenetic simulations and a phylogenetic signal based method. We also present a diagnostic analysis of the new permulations approach demonstrating how the method scales with the number of species and the number of categories included in the analysis. Our results show that our new categorical method outperforms phylogenetic simulations at identifying genes and enriched pathways significantly associated with the diet phenotype and that the new ancestral reconstruction drives an improvement in our ability to capture diet-related enriched pathways. Our categorical permulations were able to account for non-uniform null distributions and correct for non-independence in gene rank during pathway enrichment analysis. The categorical expansion to RERconverge will provide a strong foundation for applying the comparative method to categorical traits on larger data sets with more species and more complex trait evolution.

## Introduction

Determining the genomic elements underlying complex phenotypes is a fundamental question in biology. One way to address that question is through the lens of convergent evolution, or the process in which multiple species independently acquire similar phenotypes. These independent evolutionary events are natural replicates of phenotype acquisition in which similar genetic changes in correspondence with similar phenotypic changes may indicate a genotype-phenotype relationship. (Kowalczyk et al. 2019; Li et al. 2010; Partha et al. 2017). Changes in selective pressure can frequently be observed as changes in the rate of substitutions along the branches of a phylogeny due to increased constraint, relaxation of constraint, or positive selection (Yang 2007; Pond, Frost, and Muse 2005). There are a variety of methods applied to detect convergent evolution at the same individual nucleotides (Hasselmann et al. 2008), individual amino acids (Rey et al. 2018; Thomas and Hahn 2015; Fukushima and Pollock 2023), regulatory elements (Partha et al. 2017; Kowalczyk, Chikina, and Clark 2022; Kaplow et al. 2023; Yan et al. 2023), and genes (Kosakovsky Pond et al. 2020; Kowalczyk et al. 2019). These methods have been applied to identify convergent molecular evolution associated with a wide variety of traits in a wide number of clades (Yusuf et al. 2023; Jin et al. 2023).

A limitation to most comparative phylogenetic methods is the inability to analyze non-ordinal, multicategorical traits. However, some methods do exist to specifically study categorical data. Garland et al. 1993 (Garland et al. 1993) introduced the phylogenetic simulations method which uses an ANOVA test to correlate a continuous phenotype with categories and computes p-values empirically from simulations to account for phylogenetic relationships between species. Multiple phylogenetic packages including phytools (Revell 2012) and geiger (Pennell et al. 2014) have implemented this phylogenetic ANOVA test. ANOVA is a parametric test, with much stricter assumptions than the non-parametric alternatives that phylogenetic data often does not meet. Also, the method is technically designed to study *continuous* traits among species grouped into certain categories.

Departing from evolutionary rates, Ribeiro and Borges (Borges et al. 2019) presented the delta statistic for calculating phylogenetic signal for categorical traits. One of many applications of the delta statistic is identifying genetic elements associated with a categorical phenotype. It has been successfully applied to find genes associated with the evolution of nocturnality (Borges et al. 2019). However, we still note a few limitations to this method. Currently, the only option is to calculate p-values by random permutation of the trait vector. Random permutation represents the null hypothesis that there is no phylogenetic signal, but is not a good representation of the null hypothesis that a genetic element is not associated with a phenotype since we expect even null phenotypes to maintain phylogenetic relationships (Saputra et al. 2021). Additionally, the delta statistic does not take into account or provide any information on evolutionary rates, so it cannot be used in the specific context where our goal is to find genetic elements that are convergently accelerated or conserved.

To address the current limitations of RERconverge and complementary methods for categorical trait analysis, we introduce categorical methods within RERconverge, an R package for the genome-wide association of convergent evolutionary rate shifts with convergent phenotypes (Kowalczyk et al. 2019). The number of substitutions along a branch is quantified as an evolutionary rate and, correcting for certain non-specific factors, we can compute relative evolutionary rates (RERs) that reflect whether a given gene is evolving slower or faster than the genome-wide average along a branch of the phylogeny (Kowalczyk et al. 2019). We can thus use convergent evolution as a useful tool in comparative genomic analyses to identify regions of the genome showing convergent rate shifts in accordance with the phenotype.

Unlike many phylogenetic software packages, RERconverge infers the ancestral history of a trait to use when associating evolutionary rates with the evolution of the trait and provides a built-in method, known as permulations, for the correction of p-values for phylogenetic dependence and other sources of systematic variation leading to nonindependence in the data (Saputra et al. 2021). RERconverge has been used with great success to discover both coding and non-coding elements related to the evolution of continuous and binary traits including mammalian lifespan, hairlessness, and marine habitation (Kowalczyk, Chikina, and Clark 2022; Kowalczyk et al. 2020; Chikina, Robinson, and Clark 2016).

The categorical extension to RERconverge presented here infers the ancestral history of categorical traits, provide a permulation strategy for categorical traits to return reliable, phylogeny aware, corrected p-values, and have options for both parametric and non-parametric statistical tests to associate RERs with the phenotype including pairwise testing. As a test case, we applied these new methods to the diet phenotype. There are at least three commonly occurring convergent mammalian diets: carnivory, herbivory, and omnivory. Diet can be further divided into smaller categories including insectivores, piscivores, and many more. Excluding any one category to perform a binary analysis would significantly reduce the number of species in the analysis (**Fig. 1**). It is also unclear whether omnivores should be classified as carnivores or herbivores in a binary analysis, and the arbitrary choice could have potentially significant impacts on the results. These problems are likely to arise for any non-ordinal, multicategorical trait. Moreover, the encoding of a categorical trait as a binary trait does not scale with trait complexity. Applying our new methodology, we found many diet-related pathways enriched for genes with high phylogenetic signal, highlighting the potential of this method for genome-wide analyses.

**Figure 1.**
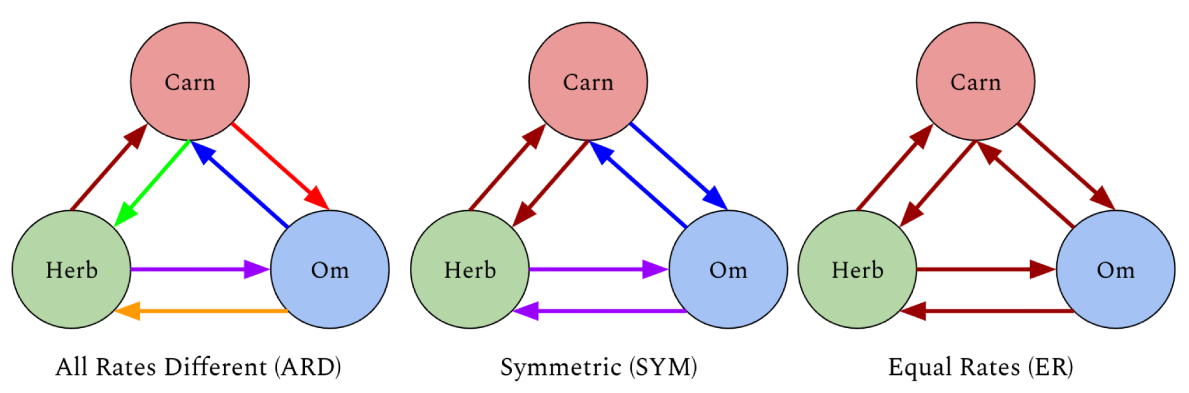
The circles represent the three categories (states) - carnivore (Carn), herbivore (Herb), omnivore (Om). The colored arrows represent the transition rates where arrows of the same color represent transitions with the same rate.

## Results

### Diet Results

#### Diet Phenotype Ancestral Reconstruction

To test our extension of the RERconverge method to categorical traits, we used the phenotype of diet (carnivore, omnivore, and herbivore) across a multiple sequence alignment of 115 high quality mammalian genomes (Hecker and Hiller 2020). The diet phenotype was annotated using prior literature across those genomes (Nowak 1999) (**Fig. 2**). The resulting annotated tree was sufficiently powered to identify diet-associated convergent evolution, with 4 direct transitions between carnivore and herbivore, 12 direct transitions between carnivore and omnivore, and 19 direct transitions between herbivore and omnivore where each transition is interpreted as a potential independent convergent event providing statistical evidence.

**Figure 2.**
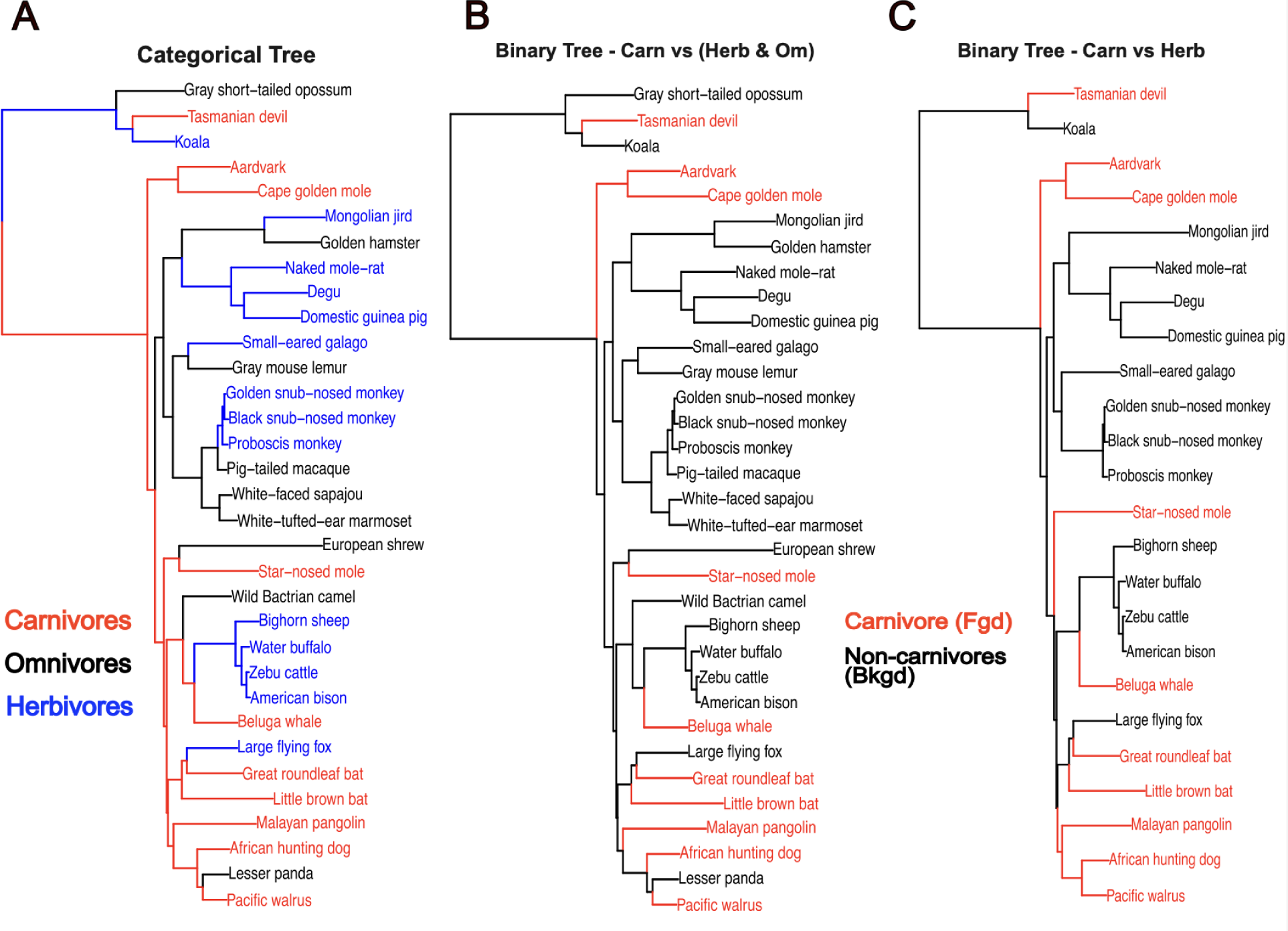
Example of a categorical reconstruction on a subset of mammals included in the full analyses (full trait reconstructions **Fig. S1**). A) Categorical reconstruction using maximum likelihood applied to a continuous time markov model. B) Binary reconstruction with carnivore foreground and herbivore/omnivore background. Uses a maximum parsimony based approach. C) Binary reconstruction on phylogeny of carnivores (foreground) and herbivores (background) with omnivores removed. Uses a maximum parsimony based approach.

We first added functionality that allowed for more sophisticated ancestral trait reconstruction that could be applied to categorical phenotypes. Previous RERconverge ancestral trait reconstruction functionality was only available for binary or continuous traits and used a maximum parsimony approach (Kowalczyk et al. 2019; Maddison 1989). Here, we added a new framework that estimates ancestral traits by applying maximum likelihood to a continuous time Markov model of evolution, which has been commonly applied to categorical traits (Pagel 1994; Pupko et al. 2000; Paradis, Claude, and Strimmer 2004).

In this framework, a rate model places constraints on the rates inferred in the transition rate matrix of the Markov model. The rate model specifies which transition rates are zero, and which rates are equal. While a user may specify any custom rate model, there are three rate models that are frequently considered. These are equal rates (ER), symmetric (SYM), and all rates different (ARD) (Paradis, Claude, and Strimmer 2004). In the ER model, all state transitions are assumed to occur with the same rate. In the SYM model, transitions are assumed to occur with the same rate in either direction, and in the ARD model, all transitions are assumed to potentially occur with a different rate (**Fig. 1**). Thus the ER model is the simplest, with only one transition rate to infer, while the ARD model is the most complex, requiring the most transition rates to be inferred. The rate model can have a significant impact on the results of the ancestral reconstruction, thus we compared these three rate models using the likelihood ratio test implemented in RERconverge (Kowalczyk et al. 2019; Pagel 1994). The ARD model provided a significantly better fit to the data (p = 0.00952 compared to ER, p = 0.02354 compared to SYM), thus was used to infer the ancestral states. Under the ARD model, the ancestral mammal was inferred to be a carnivore, while omnivores and herbivores were inferred to have evolved independently multiple times. The ancestral reconstruction inferred loss of carnivory to omnivory then return to carnivory two times in the Philippine tarsier and Chinese tree shrew lineages. This pattern of losing then regaining a trait was not possible to infer with the maximum parsimony approach employed in the prior RERconverge release (Maddison 1989). Broadly, the ancestral trait assignments by the ARD model had greater agreement with ancestral diet phenotypes annotated by experts (Price et al. 2012) than the maximum parsimony approach.

#### Methods Comparison

As described in more detail in the methods section, we performed two types of binary RERconverge pairwise analyses (Kowalczyk et al. 2019), a categorical phylANOVA analysis using the RERs computed by RERconverge, and a delta statistic analysis (Borges et al. 2019) (**Table 1**). All of these methods were tested on the same set of 19,137 gene trees and phylogeny of 115 mammals (Hecker and Hiller 2020). One important factor distinguishing these methods was the use or assignment of internal branches. PhylANOVA only includes extant species, binary RERconverge uses a maximum parsimony approach to infer ancestral states, and based on our analysis above, we chose to apply categorical RERconverge using maximum likelihood with a continuous time Markov model (CTMM) (Pagel 1994; Pupko et al. 2000; Paradis, Claude, and Strimmer 2004) to infer ancestral states.

**Table 1.** Comparison of the methods.

| Method | Works with discordant trees | Includes pairwise comparisons | P-values included (in package/function /GitHub available code)? | Approximate Runtime<sup>1,2</sup> (mins) |
| --- | --- | --- | --- | --- |
| Categorical RERconverge | No | Yes | Yes, parametric and permutations | RERs: 70<br>Analysis: 0.15<br>Permutations: 86<br>Total: 155 |
| Pairwise Binary RERconverge (method I) | No | N/A | Yes, parametric and permutations | RERs: 70<br>Analysis: 0.15<br>Permutations<sup>4</sup>: ~210<br>Total: ~ 280 |
| Pairwise Binary RERconverge (method II) | No | N/A | Yes, parametric and permutations | RERs: 120<br>Analysis: 0.75<br>Permutations<sup>5</sup>: ~360<br>Total: ~ 480 |
| phylANOVA + base RERconverge (phylogenetic simulations with Relative Evolutionary Rates)<sup>3</sup> | No<sup>3</sup> | Yes | Yes, simulations only | RERs: 70<br>Analysis: 114<br>Total: 184 |
| Delta statistic of phylogenetic signal | Yes | No | No | Total: 103 |

1. Includes time to run 500 permulations or 500 simulations with phylANOVA. Note: for this paper we ran 10,000 permulations/simulations. Does not include the time to perform a pathway enrichment analysis. Does not include the time to estimate trees from the MSA since this is universal to all the methods. The runtime for pairwise binary RERconverge method I was determined using herbivores as foreground and the runtime for pairwise binary RERconverge method II was determined using carnivores/herbivores with carnivore foreground. Note that runtimes for permulations can vary depending on the specific structure of the phenotype.
2. Runtime is for 19,137 genes and a phylogeny with 115 species
3. The base RERconverge package was used to estimate relative evolutionary rates, these were then associated with the phenotype data using phylANOVA. While phylANOVA itself works with discordant trees, computing relative evolutionary rates with RERconverge does not work with discordant trees.
4. The runtime was 70 minutes for the binary phenotype with herbivore foreground. To determine the total runtime for all three tests (one for each category as foreground), this was multiplied by 3 to obtain ∼210 minutes. This is approximate since each run of permulations varies depending on the structure of the phenotype.
5. The runtime was 60 minutes for the binary phenotype with carnivore foreground and herbivore background (omnivores removed). To determine the total runtime for all six tests, this was multiplied by 6 to obtain ∼360 minutes. This is approximate since each run of permulations varies depending on the structure of the phenotype.

The inference of ancestral states with the maximum parsimony approach used for binary traits is heavily dependent on the choice of foreground since this method steps back through the tree and assigns ancestral branches to the foreground until all the descendants are no longer part of the foreground. As a result, this approach is also sensitive to excluding species. The impact of removing omnivores on the assignment of branches to the foreground is illustrated on a subset of the full tree (**Fig. 2B** vs. **Fig. 2C**). When omnivores are removed, the common ancestor of the African hunting dog and Pacific walrus is assigned to the carnivore foreground. However, when omnivores are included, this ancestor is not assigned to the carnivore foreground because the lesser panda is an omnivore and thus not part of the foreground. The categorical reconstruction on the full tree predicts this ancestor is a carnivore (**Fig. S1**), but regardless of the right classification, we take away from this observation the sensitivity of the binary reconstruction to species presence or absence. The CTMM ancestral reconstruction is not dependent on a choice of foreground, as the ancestral state(s) versus the convergent state(s) are effectively inferred by the maximum likelihood assignment of states. Categorical reconstruction infers a carnivorous mammalian ancestor while the binary reconstructions do not infer as many carnivorous internal branches (**Fig. 2A, Fig. S1**). The exact extent of the effect of different inferred ancestral states on the gene and pathway enrichment results is most likely phenotype and phylogeny specific.

#### Comparison of gene results across methods

Association statistics between the evolution of the diet phenotype and RERs were computed for the three methods that use relative evolutionary rates: categorical RERconverge, pairwise binary RERconverge, and phylANOVA with RERs computed from RERconverge. The delta statistic method did not provide an approach for computing p-values so it was excluded from these analysis-wide gene result comparisons. 10,000 permulations were used to obtain permulation p-values for the binary and categorical RERconverge analyses. Similarly, 10,000 simulations were used to obtain simulation p-values for the phylANOVA analysis.

Quantile-quantile plots of the permulation and simulation p-values were made to determine which comparisons are driving the identification of significant genes and assess the signal across the different methods (**Fig. 3A-D**). PhylANOVA was significantly underpowered, with far fewer significant p-values in the QQ plots compared to all other methods (**Fig. 3A-D, gray**), although the same number of permulations and simulations were performed (**Fig. 3A-D, gray)**. This lack of power in PhyloANOVA could be due to lack of information on internal branches, an important feature included in RERconverge. In addition, it may also be due to differences in the phylogenetic simulations. In the ANOVA approach, RERs are treated as a continuously valued phenotype and simulated with a Brownian Motion model (Garland et al. 1993; Revell 2012). In contrast, the RERconverge approach allows for the use of phylogenetic permulations (Saputra et al. 2021), which have been shown to better correct for the tree structure.The categorical RERconverge update, which features methods to include internal branches and perform permulations, offers a significant improvement upon existing methods for comparing relative evolutionary rates across categories of species.

**Figure 3.**
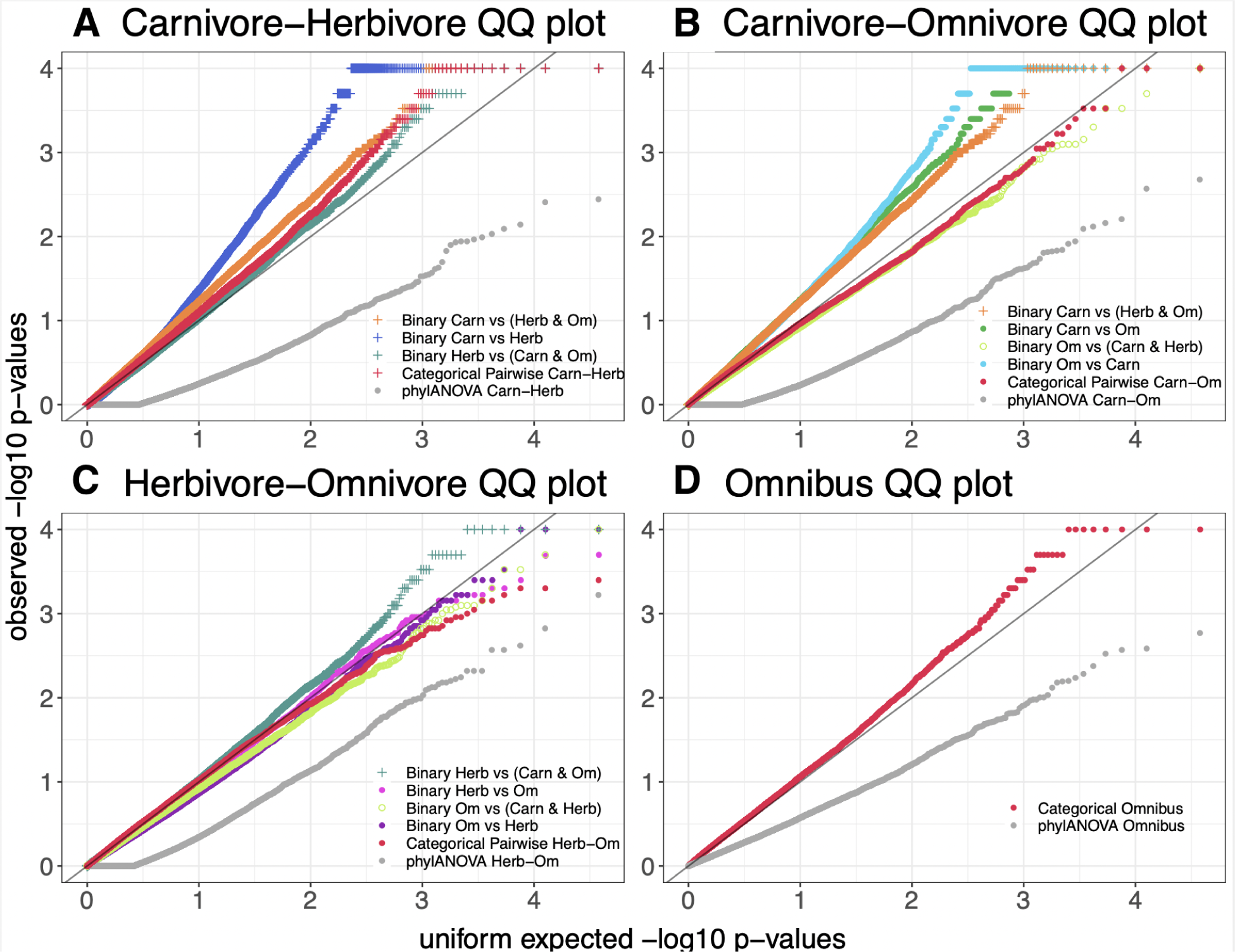
Quantile quantile (QQ) plots of the permulation or simulation p-values for the results of categorical RERconverge, phylANOVA, and pairwise binary RERconverge. A-C) A + symbol is used to denote any comparison in which carnivores and herbivores are separated between the foreground and background. C) An open circle symbol denotes the analysis in which carnivores and herbivores together form the background and omnivores form the foreground.

The binary and pairwise comparisons that measure differences between carnivores and herbivores tend to have the greatest signal, suggesting genetic differences between carnivores and herbivores are driving the results (**Fig. 3A-C, indicated by +**). For instance, among the herbivore and omnivore comparisons, the binary comparison between herbivores in the foreground and carnivores/omnivores in the background has the strongest signal (**Fig. 3C, indicated +)**. In contrast, both binary comparisons between herbivores and omnivores with carnivores removed do not show strong signal (**Fig. 3C, pink and purple)**. Furthermore, the comparison between omnivores in the foreground and carnivores/herbivores in the background does not exhibit strong signal (**Fig. 3C, open green circle**). Thus, the key feature related to strong signal appears to be the separation of carnivores and herbivores between the foreground and background.

However, the exception to the trend of carnivore vs. herbivore as the most significant comparison is within the carnivore-omnivore signal comparison, where carnivore vs. omnivore have the most signal (**Fig. 3B**).The curves for these comparisons depart from the unity line early on, suggesting that the greater signal may be the result of binary permulations under-correcting for false positives due to failing to fully correct for the phylogenetic structure in the data rather than these methods being more powerful. The effect of removing all species of one category is often to have more full clades inferred together as foreground with fewer convergent gains or losses of the trait (**Fig. 2**). This makes it more difficult to correct for phylogenetic relatedness driving the signal, especially since we used a relaxed version of binary permulations that does not enforce the permulated trees to exactly match the original phenotype tree in the structure of the relationships between foreground branches. Such differences in the inferred ancestral states due to removing herbivores may be contributing to a deceptively high degree of signal among these methods in other ways as well. These findings are consistent with our observations that internal branch assignments can have a large impact on genes’ significance.

To further explore the impact of internal branches, we focus on RERs of two genes evolving significantly slower or faster among carnivores than omnivores according to the binary methods, but showing no significant rate shift according to the categorical method (**Fig. 4**). In both cases, whether a rate shift is observed or not is driven by differences in the RER distributions of the internal branches. MADCAM1 is significantly accelerated in omnivores according to the binary analysis with carnivore foreground (p = 4.987 × 10^−7^, adjusted p = 0.0005558, permulation p = 0.0067), however this is not the case in the categorical pairwise test (p = 0.16, adjusted p = 1, permulation p = 0.2161). Separating the relative evolutionary rates by extant vs. internal species, we see that in the categorical reconstruction, the RERs of carnivores are driven upwards due to the presence of internal branches with larger RERs assigned as carnivores (**Fig. 4**). PHB2 is significantly accelerated in carnivores according to the binary analysis with omnivore foreground (p = 9.226 × 10^−5^, adjusted p = 0.01139, permulation p-value = 0.0) but not in the categorical pairwise test (p = 0.3461, adjusted p = 1, permulation p = 0.0776). The signal for this gene in the binary analysis is being driven by a group of internal branches with negative RERs. These internal branches are not assigned as omnivores in the categorical reconstruction. When this group is no longer present, there is no significant difference in RERs between carnivores and omnivores (**Fig. 4**).

**Figure 4.**
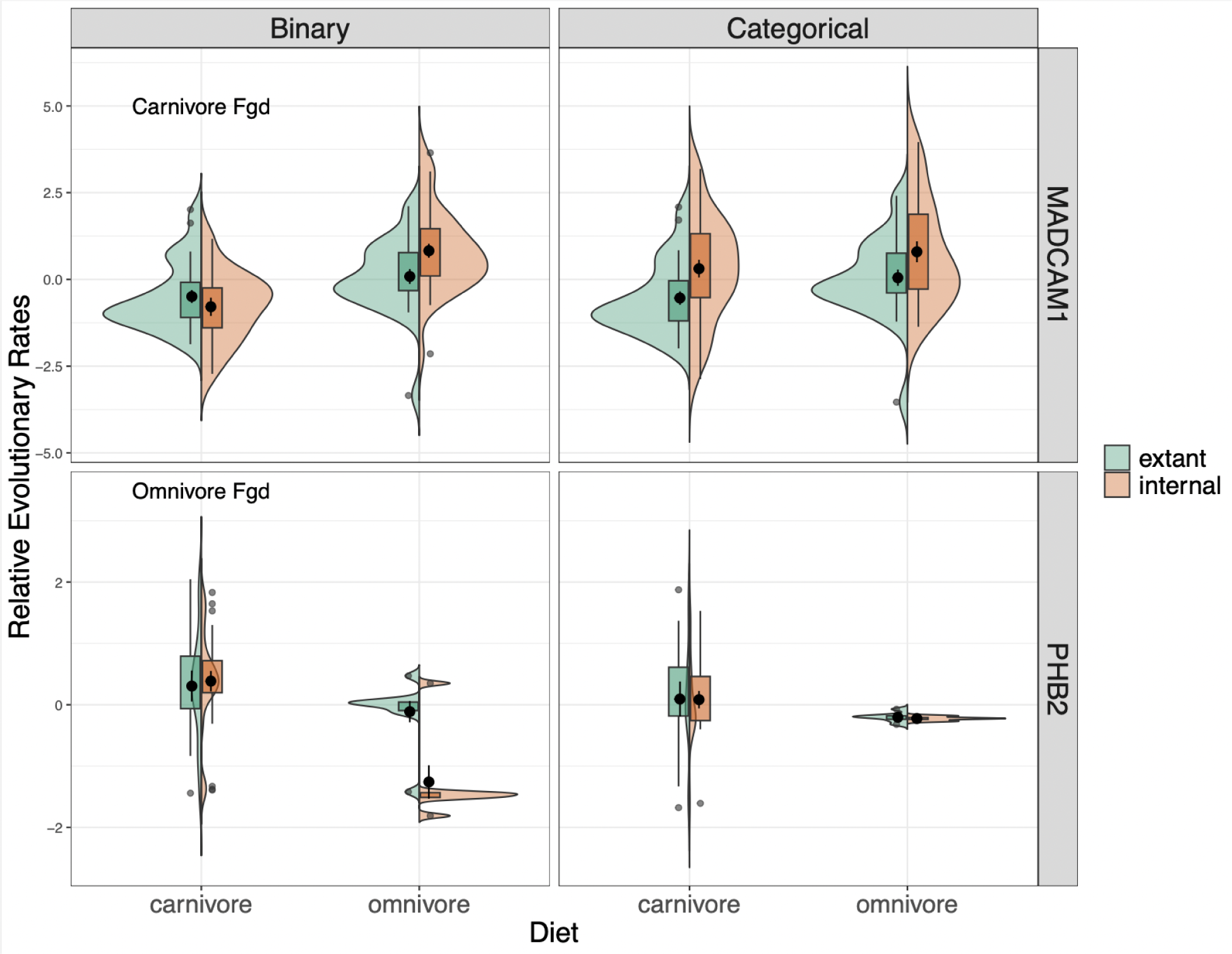
Each violin plot compares the distribution of RERs between carnivores and omnivores for MADCAM1 (top row) and PHB2 (bottom row). In green, on the left, is the distribution of RERs for extant branches and in orange, on the right, is the distribution of RERs for internal branches. The left column shows the RERs from the binary analyses of carnivores vs. omnivores with either carnivore foreground (top) or omnivore foreground (bottom). The right column contains the RERs from the categorical analysis.

#### Comparison of pathway enrichment results across methods

Effects of the ancestral reconstruction extend beyond the gene level results to the pathway enrichment results as well. One GO pathway for which we may expect to see significant enrichment is digestive tract development. Carnivores have been shown to have shortened digestive tracts (Kim et al. 2016) and we would expect the digestive tract to specialize to a species’ diet. Digestive tract development was significantly enriched in the categorical pairwise test between carnivores and herbivores (adjusted p = 0.044, permulation p = 0.0011) (**Fig. 5A, blue line**). The four different binary pairwise analyses between carnivores and herbivores are also shown. Enrichment is lower among these four methods compared to the categorical method. Notably, the two methods with herbivore foreground (**Fig. 5A, purple and green**) have greater enrichment than the method with carnivore foreground. This is consistent with the categorical ancestral reconstruction inferring more carnivorous ancestral species than herbivores (**Fig. S1**) so these binary ancestral reconstructions are more similar to the categorical ancestral reconstruction. This suggested that the ancestral reconstruction was playing an important part in the enrichment of this pathway.

**Figure 5.**
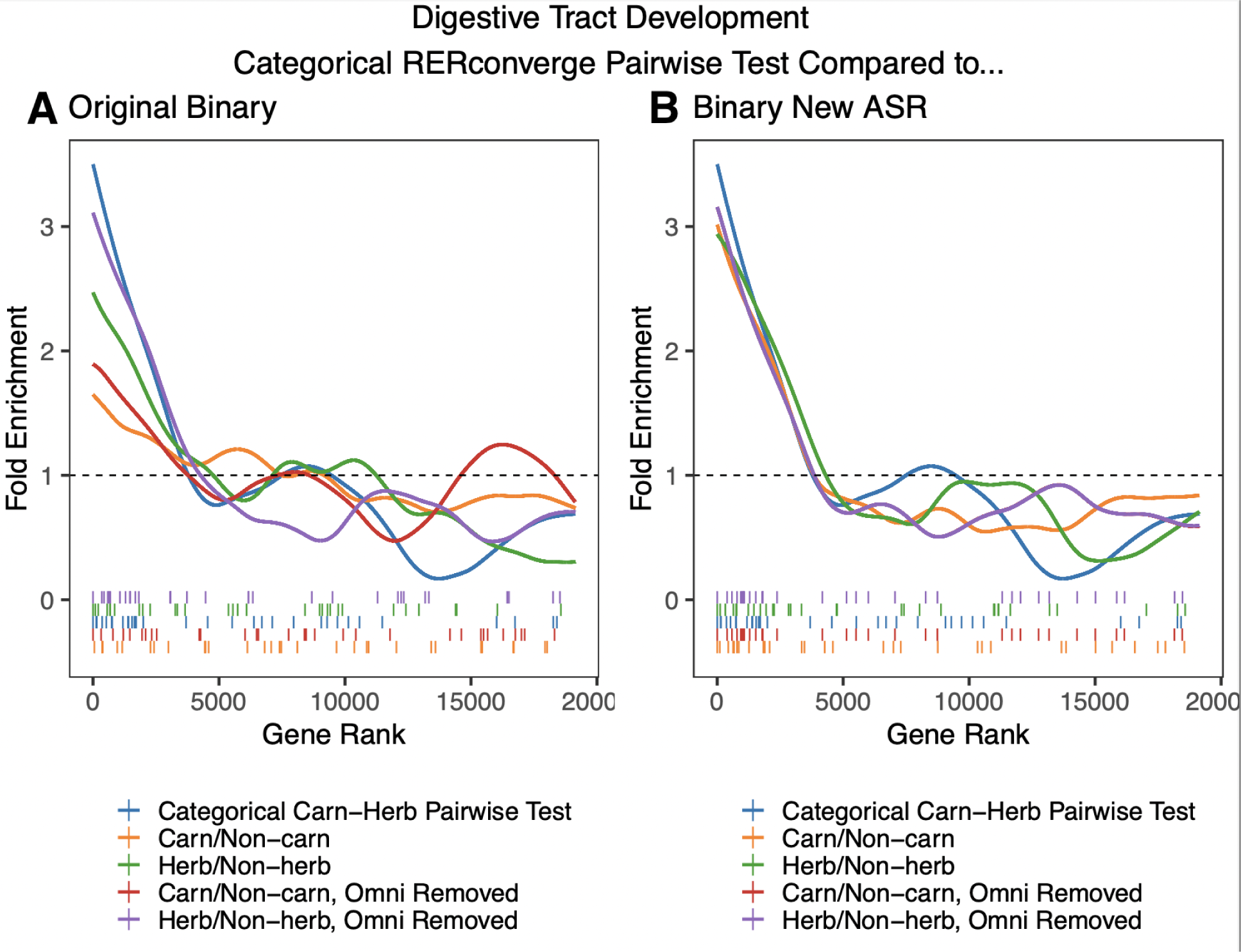
Fold enrichment and barcode plots for the digestive tract development pathway. A) The categorical carnivore-herbivore pairwise test (blue) is compared to the four different original binary analyses which compared carnivores to herbivores. B) The categorical carnivore-herbivore pairwise test (blue) is compared to the four binary RERconverge analyses in which the new ancestral state reconstruction (ASR) method, maximum likelihood applied to a continuous time Markov model, has been used in place of the original maximum parsimony approach.

To specifically test the effect of the ancestral reconstruction method relative to different statistical tests employed, we used the CTMC ancestral reconstruction method to infer the ancestral states for each of the four binary phenotypes. We then performed the rest of the binary analysis in the same way as before (**Fig. 5B**). Note that the two binary methods in which omnivores were removed (red and purple lines) now had the same assignment of ancestral states because the reconstruction is no longer dependent on foreground choice. Using the new ancestral reconstruction caused the enrichment of digestive tract development genes to improve, reaching nearly the same level as the categorical pairwise test. Thus, we infer the differences in enrichment were being driven almost entirely by the ancestral reconstruction. Using the more sophisticated ancestral reconstruction resulted in improved enrichment for a pathway which we would expect to be enriched, suggesting that capturing more complex patterns of evolution and using more reliable reconstructions is beneficial at the pathway enrichment level as well. The new ancestral reconstruction also appears more robust to removing species from the analysis because in the results from the new reconstruction, the red and purple lines, in which omnivores were removed, are much more similar to the orange and green lines respectively, in which omnivores were not removed, compared to the old reconstruction (**Fig. 5B** vs. **Fig. 5A**)).

Pathway enrichments for sets of MGI and GO pathways (Subramanian and Others 2005; Liberzon et al. 2011) were computed for all binary RERconverge analyses, the categorical RERconverge analysis, the phylANOVA analysis, and the delta statistic analysis. For the binary and categorical RERconverge analyses, the genes were ranked by their parametric p-values. Permulations were then performed at the pathway level to correct both for non-independence among the species and for non-independence in gene rank (groups of genes in pathways often shift together in rank) (Saputra et al. 2021). The gene association results from the 10,000 permulations were used to recalculate pathway enrichments for each permulation, providing 10,000 null enrichment statistics for each pathway. The permulation p-value for a pathway is the proportion of null statistics more extreme than the observed enrichment statistic.

The distributions of enrichment permulation p-values for the RERconverge analyses did not show much signal with the exceptions of the binary omnivore vs. carnivore and carnivore vs. herbivore analyses (**Fig. 6A**). However, using the carnivore/herbivore enrichment results of MGI pathways as an example, we demonstrate that greater signal does not necessarily appear to correspond with more refined diet-related results among the RERconverge analyses. This is illustrated as the proportion of digestive system, liver/biliary system, or metabolism related pathways out of the total number of pathways identified by that method as determined by meeting an adjusted p-value threshold of 0.15 and, for the RERconverge methods for which permulations were performed, a permulation p-value threshold of 0.025 (**Fig. 6B**). The phylANOVA analysis, on the other hand, does show lower power to detect these diet-specific pathways (**Fig. 6**).

**Figure 6.**
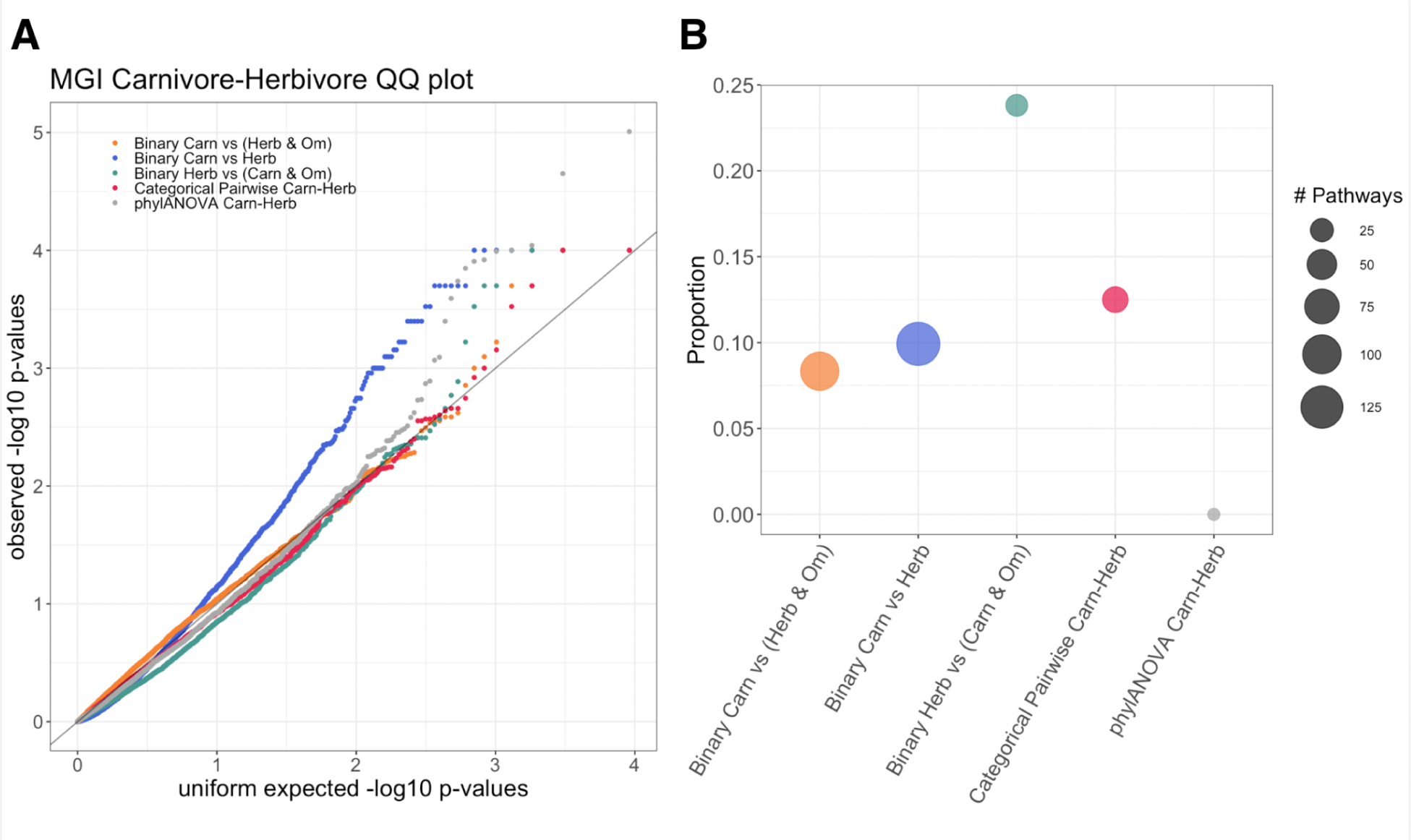
A) Quantile-quantile plot of the enrichment p-values (permulation p-values for the RERconverge results and adjusted p-values for the phylANOVA results). B) The proportion of top enriched pathways that are digestive system, liver/biliary system, or metabolism related. The size of the dots represents the total number of pathways for that method meeting the significance threshold of p.adj < 0.15 and, for the RERconverge methods, permpval < 0.025.

We showed that phylogenetic simulations, implemented by phylANOVA, did not provide the same level of statistical power as categorical RERconverge (**Fig. 6B**). Additionally, we showed that while the statistical power of categorical RERconverge is comparable to that of pairwise binary RERconverge, we are likely to see improvements from using the CTMC ancestral reconstruction methods due to the importance of inferred internal branches in the analysis. We next demonstrate how categorical RERconverge performed on a few specific pathways and genes to highlight its robust ability to identify convergent shifts in evolutionary rates and to incorporate information from multiple pairwise comparisons to identify enriched pathways.

#### Categorical RERconverge analysis results for genes and specific pathways

The differences in the ability to detect diet relevant pathways with the different methods are highlighted by the RERs of two genes within the pathway, ITGB4 and ITGA6 (**Fig. 7**). They encode integrin subunits which tend to associate to form a heterodimer (O’Leary et al. 2016), and interestingly both show acceleration among carnivores compared to herbivores (**Fig. 7A,B**; ITGB4 p = 5.662 × 10^−14^, adjusted p = 1.0719 × 10^−9^, permulation p = 0.0; ITGA6 p = 5.74 × 10^−5^, adjusted p = 0.00407, permulation p = 0.006). These genes show different patterns relative to omnivores; ITGB4 illustrates that the distributions of relative evolutionary rates may differ significantly between all three categories while ITGA6 is not significantly different between carnivores and omnivores.

**Figure 7.**
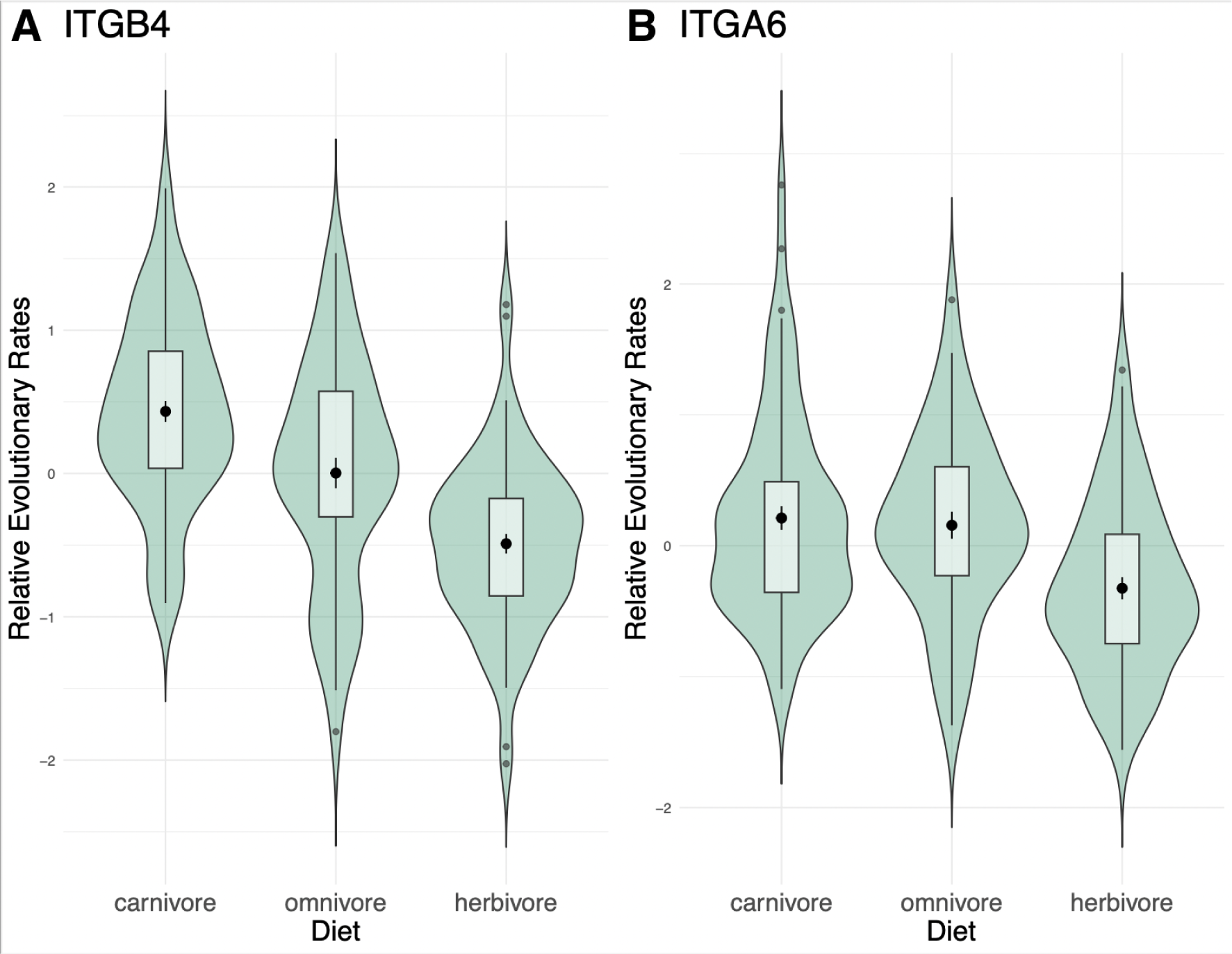
Violin plots of the distribution of relative evolutionary rates across each diet category for A) ITGB4 and B) ITGA6.

Having interpreted the updated RERconverge results, we next conducted a comparison of our method to the delta statistic. Based on its clear diet relevance and strong enrichment, we conducted the comparison on the genes within the abnormal digestion pathway (**Fig. 6**). We found significant enrichment of the categorical omnibus test and the delta statistic results but none of the pairwise or binary analyses (**Fig. 8, blue, orange;** categorical omnibus p = 6.668 × 10^−5^, adjusted p = 0.1015, permulation p = 9 × 10^−4^; delta statistic p = 0.0002565, adjusted p = 0.05586). This lack of enrichment remained even when performed with the CTMC ancestral reconstruction as was done with the digestive tract development pathway (**Fig. 5**). This suggests that the categorical test is capturing effects across multiple pairwise comparisons which alone aren’t sufficient to detect enrichment for this pathway.

**Figure 8.**
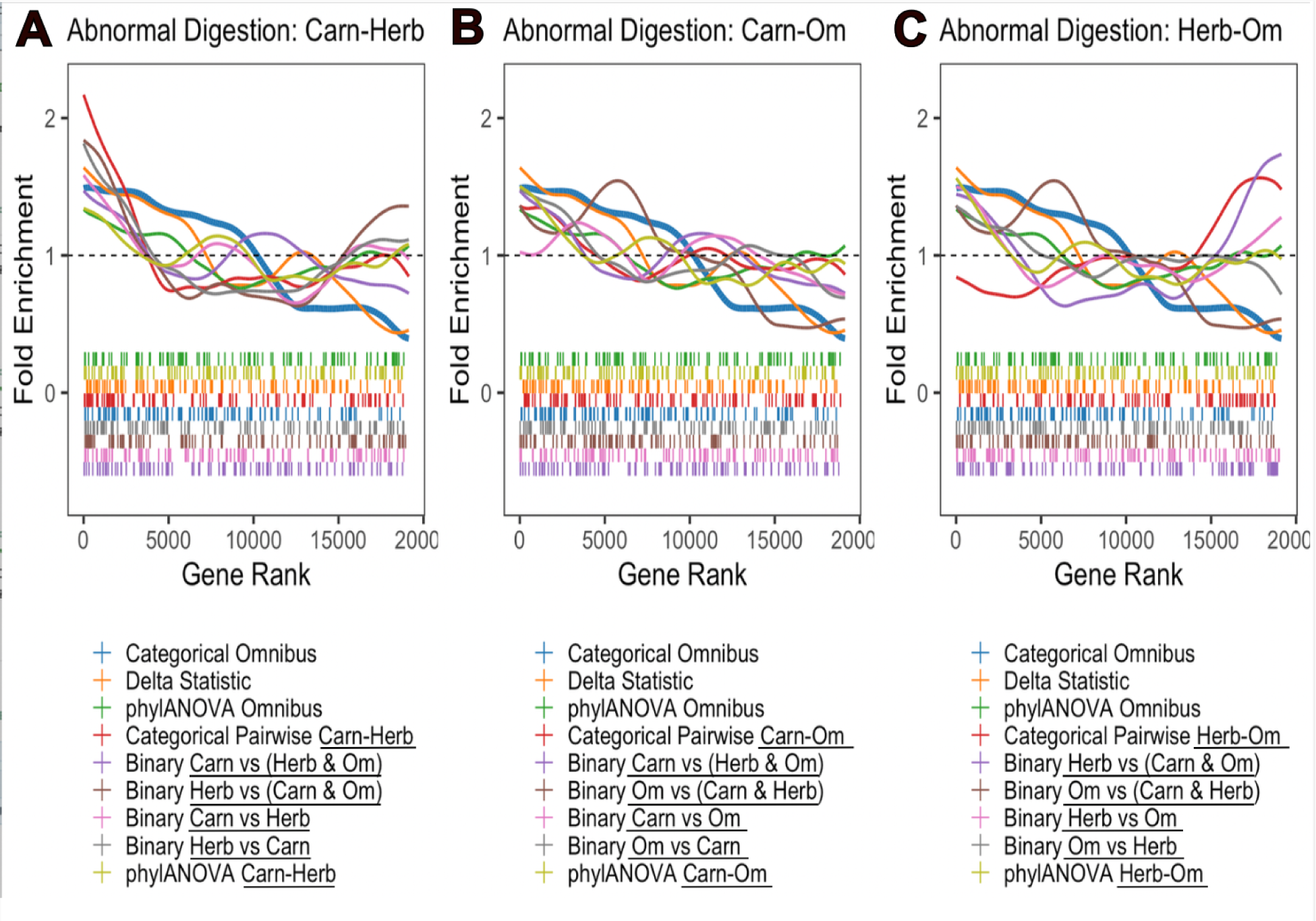
Fold enrichment and barcode plots for the abnormal digestion pathway. All three plots contain the categorical omnibus curve (blue, bold), delta statistic curve (orange), and phylANOVA omnibus curve (green). A) In addition to the omnibus methods, the methods comparing carnivores to herbivores are included. B) In addition to the omnibus methods, the methods comparing carnivores to omnivores are included. C) In addition to the omnibus methods, the methods comparing herbivores to omnivores are included. In A-C, the same color is used for *corresponding* methods. Where the corresponding methods may differ in which specific categories are being compared, the category names are underlined in the legend.

To interpret the improved performance of the omnibus statistic, we next explored the behaviors of the top 31 genes. There are two important factors which may be contributing to the stronger enrichment under the omnibus test compared to the pairwise tests. First, top genes within this pathway exhibit significant rate shifts in opposite directions under the same pairwise comparison(s). For instance ITGB1 is evolving slower in carnivores compared to herbivores and omnivores whereas TREH is evolving faster in carnivores compared to herbivores and omnivores (**Fig. 9**). RERconverge determines enrichment using a wilcoxon rank sum test to identify whether genes within a pathway are together shifted towards accelerated evolutionary rates or towards slower evolutionary rates. As a result, genes within the same pathway with evolutionary rate shifts in opposite directions do not both contribute to the same enrichment signal and may even “cancel out” each other’s effect. The omnibus test is one-sided, so significant genes move towards higher ranks together regardless of the direction of the rate shift. Second, significant differences in evolutionary rates for genes in this pathway arise among all three pairwise comparisons, not just one. Though globally the carnivore-herbivore pairwise comparison had the greatest signal (**Fig. 3A**), there are multiple top genes within this pathway whose individual signal is being driven by differences in evolutionary rates between carnivores/omnivores and/or herbivores/omnivores instead (**Fig. 9, indicated by black boxes)**. These were the two overall less powerful comparisons in the categorical analysis (**Fig. 3B-C**), however this emphasizes that important information would be lost by excluding them.

**Figure 9.**
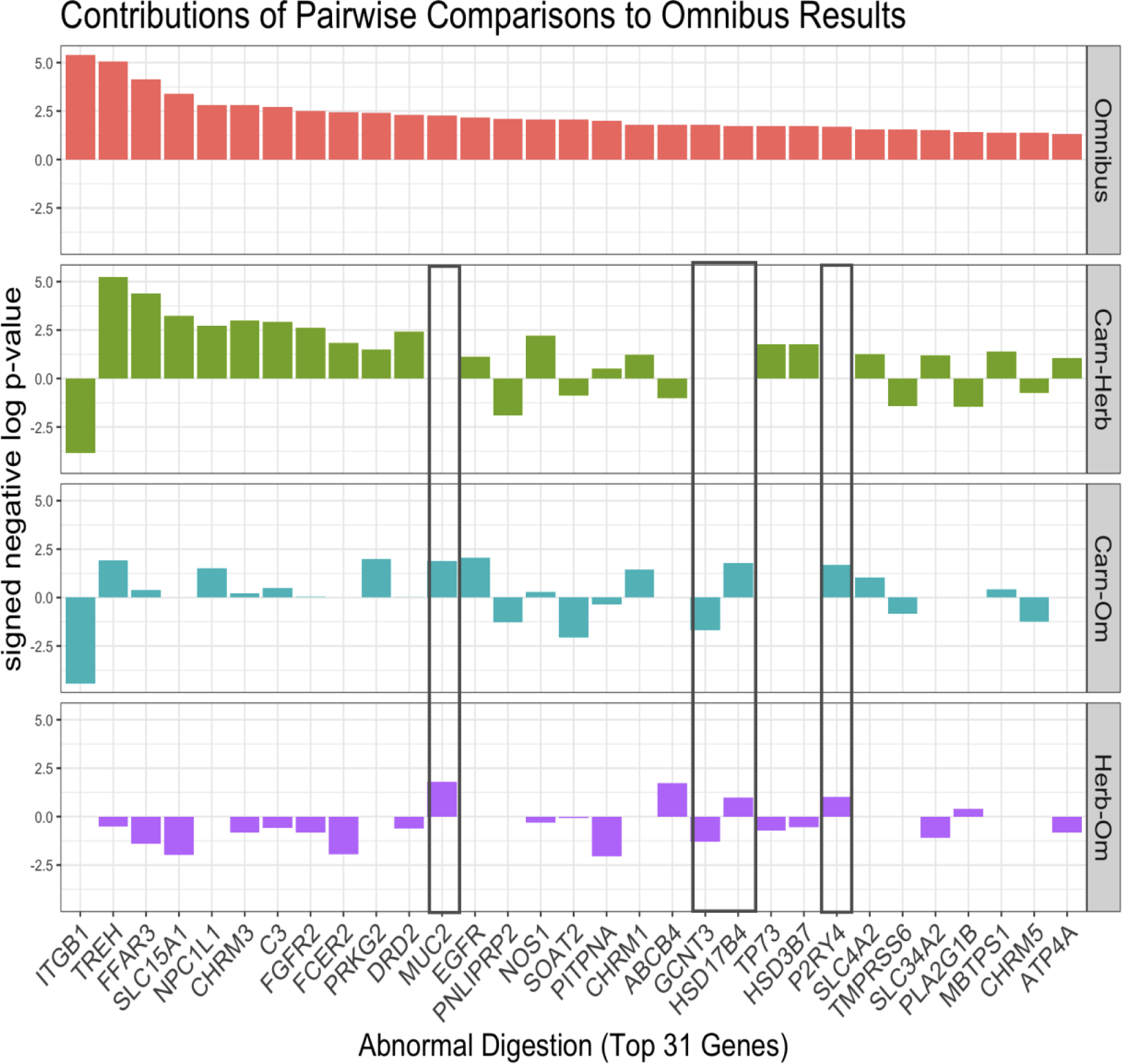
Bar plots of the top 31 genes in the abnormal digestion pathway (as ranked by the omnibus results) for the categorical omnibus and pairwise tests. The height of the bar represents the signed negative log p-value.

The delta statistic method is markedly different from the other methods in this analysis in that it does not use evolutionary rates. More details on how the delta statistic is calculated are given in the delta statistic paper (Borges et al. 2019; Ribeiro et al. 2023) and summarized in the methods section. In order to determine if the delta statistic and categorical RERconverge identified similar or different signatures of convergent evolution, we first compared both methods on the abnormal digestion pathway, which was significantly enriched according to both. The most significant genes in the abnormal digestion pathway identified by the delta statistic were for the most part different from those identified by omnibus categorical RERconverge (**Fig. 10A**). Though there’s no clear relationship between the delta statistic and the omnibus results, patterns emerge when the results are broken down into their pairwise comparisons. The most significant delta statistic genes tended to be those which were evolving slower in carnivores compared to omnivores or herbivores as indicated by negative statistics (**Fig. 10A**). Genes in the pathway which were ranked least significant according to the delta statistic, but which exhibited a rate shift, tended to be those evolving faster in carnivores compared to herbivores or omnivores (**Fig. 10A**).

**Figure 10.**
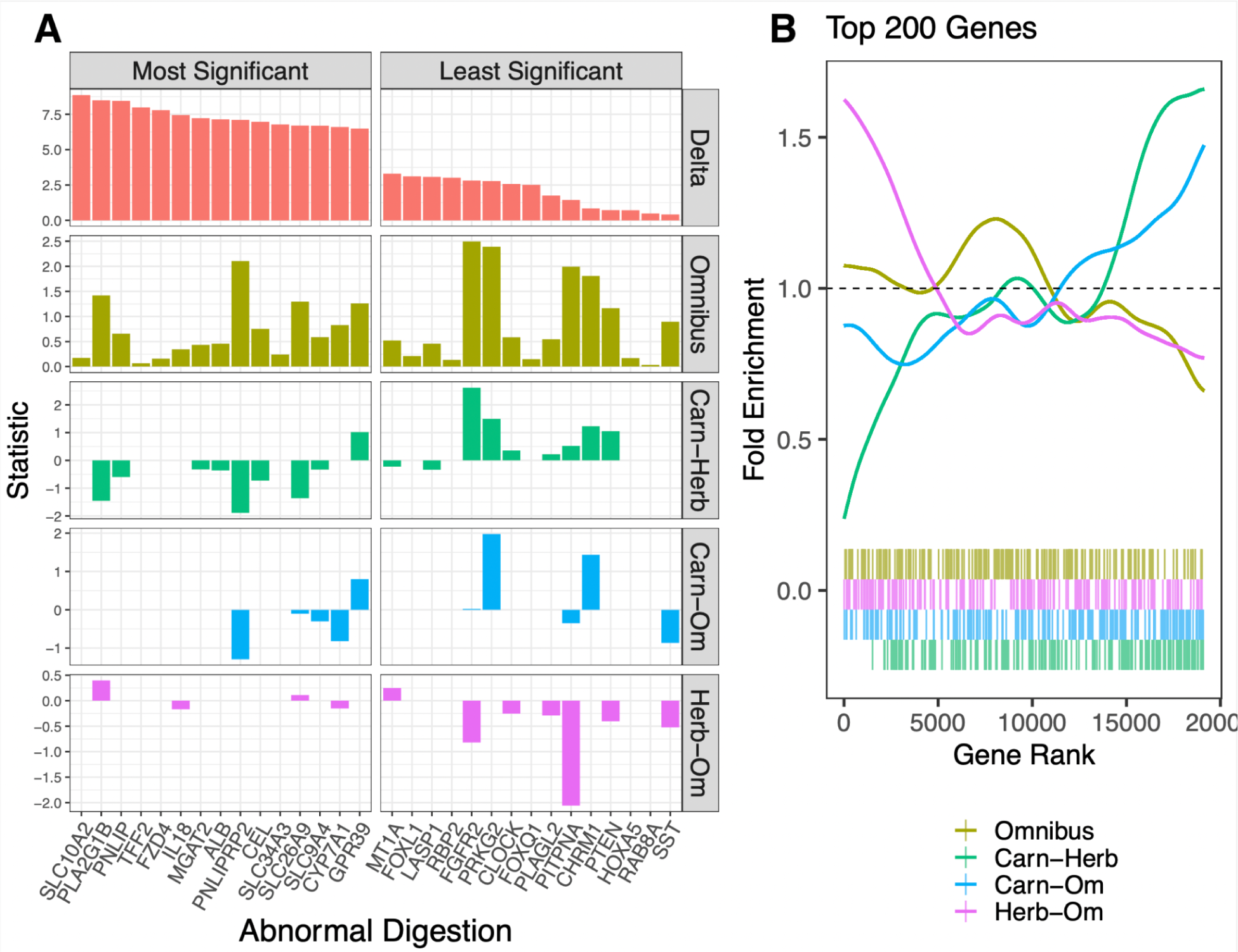
A) Bar plots of the statistic used to calculate enrichments for the top 15 most signficant genes in the abnormal digestion pathway (right) and 15 least signficant genes in the abnormal digestion pathway (left) as ranked by the delta statistic. For the delta statistic method, the statistic is delta, the measure of phylogenetic signal. For the RERconverge methods, the statistic is the signed log p-value. B) The fold enrichment and barcode plots among the RERconverge omnibus and pairwise tests of the top 200 genes ranked by the delta statistic of phylogenetic signal.

To determine if this trend persists across all genes in the analysis, not just those within the abnormal digestion pathway, we plotted the fold enrichment of the top 200 delta statistic genes among the categorical RERconverge tests (**Fig. 10B**). The general results indeed recapitulate what we observed within the abnormal digestion pathway. Genes with greater phylogenetic signal for the diet phenotype tend to evolve slower in carnivores compared to herbivores (**Fig. 10B, green)**. To a lesser extent, the same trend is observed for carnivores compared to omnivores (**Fig. 10B, blue**) and herbivores compared to omnivores (**Fig. 10B, pink**). There is no enrichment of the top delta statistic genes among the omnibus results (**Fig. 10B, yellow**), indicating that overall the two methods are identifying different, though potentially complementary, top genes associated with this diet phenotype.

This trend may be the consequence of the delta statistic using reconstruction certainty to identify phylogenetic signal. Reconstruction certainty, among many factors, depends on inferred phylogeny branch lengths which are related to evolutionary rates. We may expect a phylogeny’s ancestral history to be inferred with greater or less certainty when the branch lengths also exhibit a certain pattern as a result of allowing more or less change along the branches in order to give rise to the observed phenotype structure. In this case, slower evolutionary rates among carnivores may improve the certainty of inferring carnivorous ancestors corresponding to what we’d expect from our maximum likelihood reconstruction on the master tree (**Fig. S1**), leading to higher delta statistics. However, it is unclear to what extent these patterns reflect the real biology or methodological uncertainty in the inferred branch lengths, gene tree structure, and ancestral likelihoods.

We interpret this to mean that the delta statistic is sometimes able to recognize features distinct from those captured by the evolutionary rates in RERconverge, however that the delta statistic may be more likely to fail to detect phylogenetic signal for certain genes depending on their patterns of convergent evolutionary rate shifts.

#### Permulations Results

Previously, binary and continuous permulations was demonstrated to refine pathway enrichment results by taking nonindependence of gene ranks into account (Saputra et al. 2021). We demonstrate that categorical permulations has the same ability to refine pathway enrichment results. For example, we observed that olfactory genes commonly cluster together in rank. As a result, olfactory signaling and olfactory transduction were two of the most significant pathway enrichment results according to non-permulated p-values. These had more significant p-values than the starch and sucrose metabolism pathway from the canonical gene set (Subramanian and Others 2005; Liberzon et al. 2011) which has a clear relationship to the diet phenotype. After permulations, starch and sucrose metabolism was retained as the top result (based on permulation p-values), while olfactory signaling and olfactory transduction were no longer among the top enriched pathways (**Fig. S2B**).

We also expect categorical permulations to account for non-uniform null distributions. To determine if this was the case, we plotted histograms and quantile-quantile plots of the parametric and permulation p-values (**Fig. S2A**). The categorical pairwise tests all showed a large enrichment for high parametric p-values of around 1. This enrichment of high p-values is also detected in the permulations and is thus no longer present among the permulation p-values. This demonstrates that the enrichment of high p-values is most likely specific to this statistical test and sources of nonindependence in the data rather than representing a true pattern specific to the diet phenotype.

Notably, the simulation p-values calculated by phylANOVA still have an enrichment of high p-values near one (**Fig. S2A**). In fact, this enrichment is even more extreme than that of the categorical pairwise parametric p-values. PhylANOVA simulates the RERs using a Brownian Motion model, so while this takes phylogenetic dependence into account, it doesn’t account for other systematic variation such as nucleotide content or genome quality that can affect the RER values themselves and lead to non-uniform null distributions. The enrichment of parametric and phylANOVA simulation p-values near 1 and removal of such enrichment by permulations was observed across all the pairwise tests (**Fig. S2A**). Thus, unlike permulations, phylogenetic simulations were not able to correct for this pattern.

Additionally, though the same number of simulations and permulations (10,000) were performed, the categorical RERconverge pairwise test permulation p-values show more power, which may be because phylANOVA only uses extant species while RERconverge uses ancestral species as well.

#### Permulations Timing

Categorical permulations demonstrate how we can expand phylogenetic permulations to work with categorical phenotype data, however the current implementation of this method has limitations. The rejection sampler (see methods, step 1) becomes very slow for large trees with greater than three categories, especially in phenotypes which are more highly clustered according to phylogeny. To allow for comparison between the number of categories used in the permulations, the diet phenotype was further subset into six phenotypes, (omnivore, carnivore, herbivore, insectivore, piscivore, and anthropivore). Anthropivore, while a poorly described phenotype, was included as it was used only in its capacity as a useful subset of omnivores.

We compared both the effect of species number included in the analysis and number of categories used in the analysis on the time required to complete permulations. While the number of species did not appear to have an effect on permulation time, the number of categories in the analysis had a drastic effect. We used linear regression to assess the effect of the number of species included in the analysis on the time to perform the analysis. In the two-category analysis, where species number appeared to be a major factor in permulation time (R^2^ > 0.5), the effect appeared to be minor (0.02 seconds per species) (**Fig. 11A**). For analyses with more than 4 categories, species number appeared to have no substantial effect on time (R^2^ <0.1) (**Fig. 11, B,C,D,E**). In a few cases where outliers were removed, increased species number appeared to have a reducing effect on permulation time.

**Figure 11.**
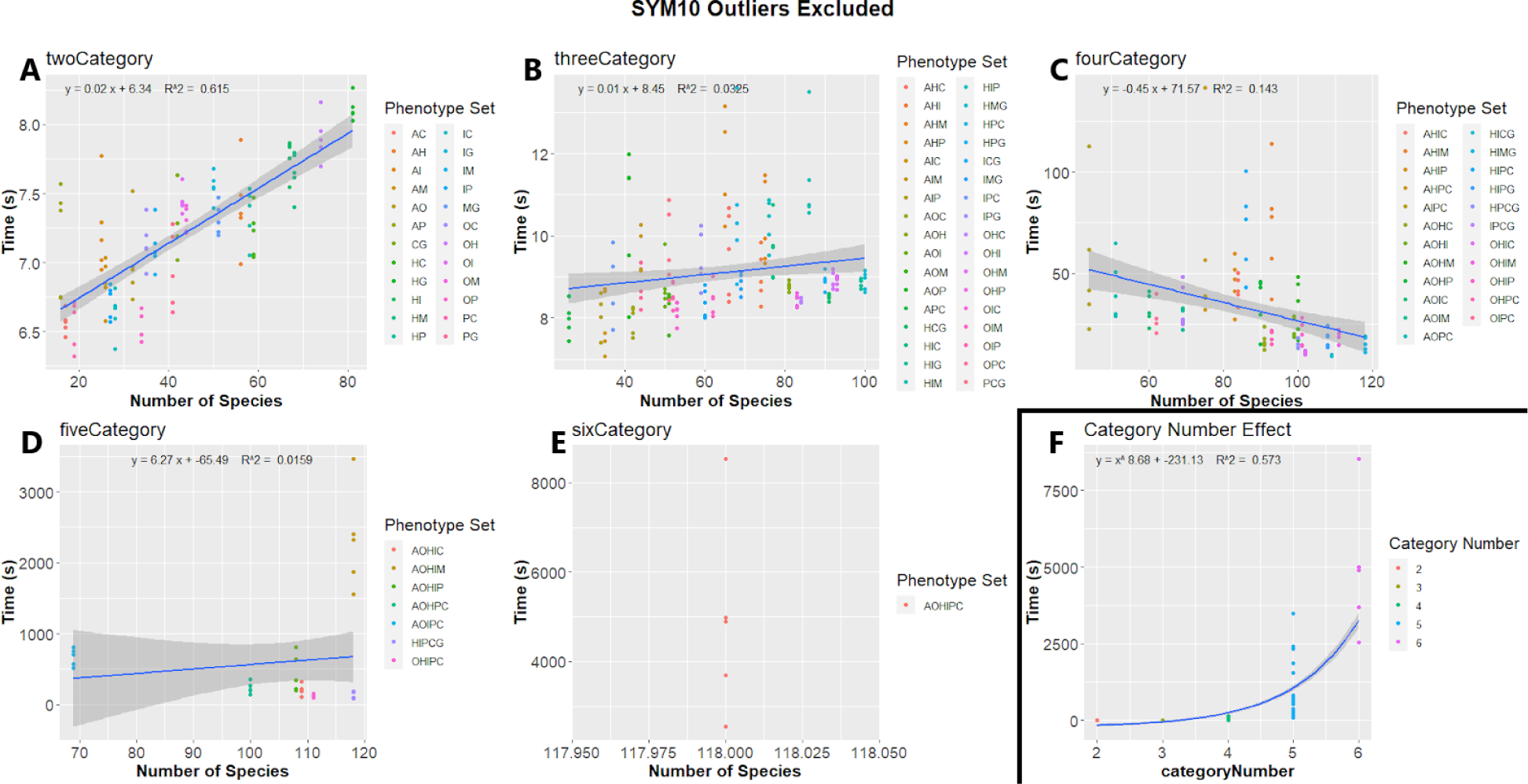
Linear regressions of time to complete permulations against number of species included in permulations. For two and three category analyses, species number has an effect, but it is minor (0.02 seconds per species). For analyses with four or more categories, there does not appear to be an effect. Note the differences in y-axis scale between category numbers. F shows the relationship between category number and permulation runtime, with each point representing the average of a full phenotype set.

We first compared the effect of category number on permulation time by comparing the average time to complete a phenotype with each number of categories. The time required increased dramatically with the number of categories. Time (seconds) per phenotype set increased exponentially (y= x#x005E;8.11 −223.66, R^2^ = 0.567) as the number of categories increased, ranging from 7.2 seconds for two categories to 4,933 seconds for six categories (**Fig. 11,F; Table S1**).

Additionally, we directly compared identical species sets with different category numbers, based on combining mergeable phenotypes. For instance, [Herbivore-Carnivore-Piscivore] was compared to [Herbivore-<Carnivore&Piscivore>]. The singly-combined phenotype sets were found to be faster than the unmerged phenotypes, particularly when there were more than 4 categories in the unmerged analysis (**Fig. 12**). On average, the merged phenotypes were 23%±14%, 284%±144%, and 1502%±762% faster at 3v2, 4v3, 5v4, respectively once outliers were removed. Two 6v5 comparisons exist in the dataset, with the merged phenotype being faster but to highly different degrees (speed increase of 112% and 3319%). The double combination phenotypes were found to be dramatically faster than the unmerged phenotypes, being 481%, 5014±3484%, and 32,220% faster at 4v2, 5v3, and 6v4 respectively. This is further evidence that the runtime increase grows more pronounced as additional categories are added. Note that these results are from a relaxation level of 10%, and as such the differences between category number is even more pronounced in unrelaxed permulations (**Table S2, Fig. S5**).

**Figure 12.**
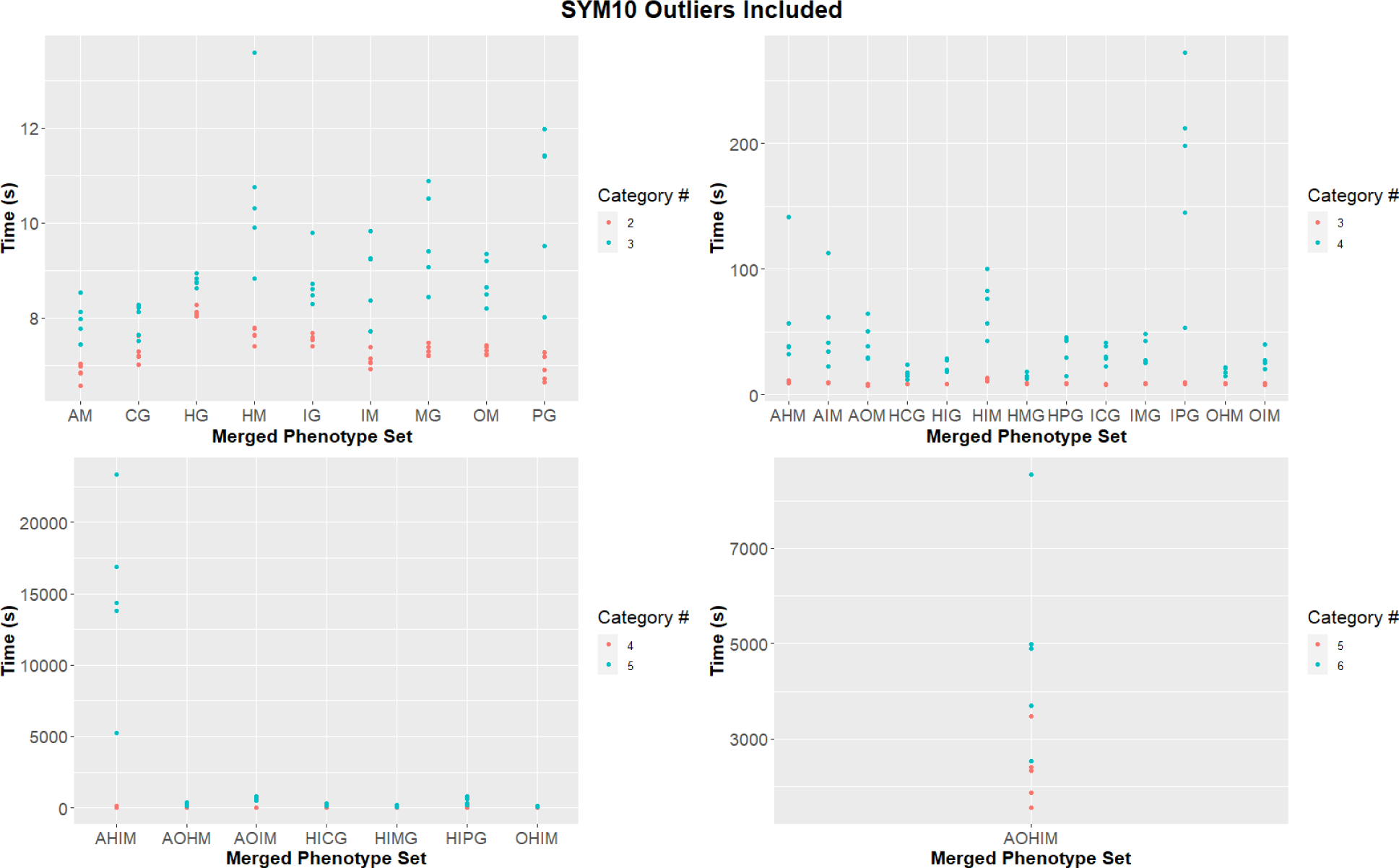
Plots of runtime in seconds of all singly-combinable phenotypes; when combined two (A), three (B), four (C), and five (D) categories, showing both merged and unmerged times. Unmerged phenotypes sets (eg. Herbivore-Carnivore-Piscivore) are shown in blue, merged phenotype sets (eg. Herbivore-<Carnivore&Piscivore>) are shown in red. The X-axis displays the first letter of the phenotypes used in the merged phenotype set.

#### Permulations Relaxation

To address these limitations, we developed a relaxed version of permulations that does not enforce such strict rejection constraints during the first step (Methods). We tested the speed increase from relaxation by running the timing test for each phenotype set at 0%, 5%, 10%, and 20% relaxation. The speed increase from relaxation is dependent on the number of categories, increasing non-linearly as the number of categories increases (**Fig. 13, Table S3**). For two categories, the improvements are marginal: with a mean improvement across all phenotype sets of 1.3%, 12%, and −1.5% for 5%, 10%, and 20% relaxation respectively. For three categories, the improvements were 3%, 57%, and 63%. For four categories, the improvements were 18%, 646%, and 1991%, dramatically faster than unrelaxed permulations. While unrelaxed versions of 5 and 6 category phenotype sets could not be completed, the 20% relaxed permulations were faster than the 10% relaxed permulations by 964% for five category sets, and by 40,000% for the six category set. When outliers were removed from the data, the speed increase was more pronounced.

**Figure 13.**
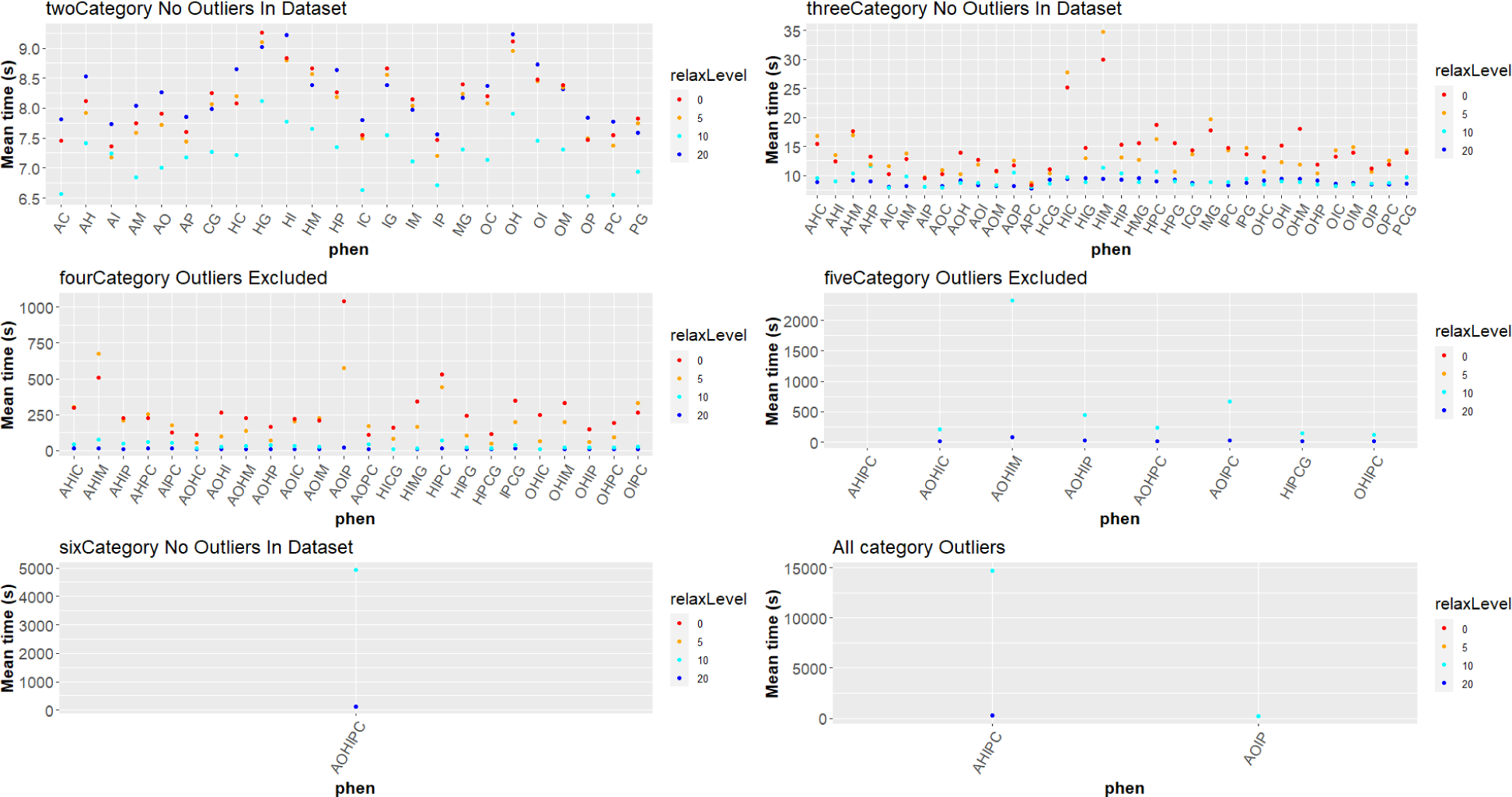
Plots of average time in seconds required to complete each phenotype set, arranged by category number (A,B,C,D,E). Note the difference in y-axis scales between categories. Outliers are present in a separate plot (F) to allow for readable scales. Each point represents the average of five time trails of the phenotype, with the color of the point indicating the relaxation level of the time trials.

The relaxed and unrelaxed permulation p-values had a mean difference of <0.03 for the omnibus and all pairwise comparisons for both IPC and HMG, and no significant difference in the effect of relaxation on p value between IPC and HMG.

In order to verify that a small relaxation of 10% on the extant species counts while maintaining internal category counts exactly as they are in the original ancestral reconstruction did not noticeably reduce the evolutionary plausibility of the permulated phenotypes, we generated 10,000 permulations and computed the log likelihood of each one (see methods, equation 1). We plotted the distributions of the log likelihoods of the simulated phenotypes that passed the rejection sampler (see methods, step 1) against the distributions of the log likelihoods of the permulated phenotypes that were generated from these simulations (see methods, steps 2 & 3). We expect plausible permulated phenotypes to have log likelihoods similar to the simulations. The relaxation did not greatly affect the distribution of log likelihoods of the permulated phenotypes (**Fig. S3**). Thus using a small relaxation can significantly speed up the permulations, but is not expected to harm the evolutionary plausibility (log likelihoods) of the permulated trees.

## Discussion

Here, we extend the RERconverge method to include the analysis of categorical traits. Our update included updated methodology for ancestral trait reconstruction, the application of statistics that can be applied to categorical variables, and the adaptation of permulations to correct for categorical traits. To benchmark these features, we used the enrichment of diet-related pathways, “digestive tract development” and “abnormal digestion” in the categorical analysis of diet traits. Our results on the “digestive tract development” pathway demonstrate the importance of improving ancestral trait reconstruction with MCMC, even when binary phenotypes are used. Our results on the “abnormal digestion” pathway demonstrate that categorical analysis can identify convergent evolution in relevant genes even when a series of binary pairwise measurements fail. Our methods have been added to the RERconverge pathways and are already freely available (Kowalczyk et al. 2019). For those intending to use our methods, if runtime is a concern, when comparing four or more categories we recommend using a relaxation level of 10% for permulations. While exact effects differ between trees, using 10% relaxation provided order of magnitude speed increases without significantly affecting results.

In addition to comparing results across different ways of running RERconverge, we compared diet pathways results to an ANOVA approach (Revell 2012) and to a delta statistic approach (Borges et al. 2019). Although the ANOVA approach has been extensively used, it performed poorly in finding diet signal relative to our RERconverge updates and the delta statistic. The delta statistic and categorical RERconverge both show diet-related signal, but genes driving the signal differed between the two methods. For example, for the “abnormal digestion” pathway, the enrichment results were similar between the categorical omnibus test we developed and the delta statistic approach. However, different genes contributed to the enrichment - ITGB1 and TREH were captured by RERconverge omnibus test while SLC10A2 and PNLIP showed greater enrichment in the delta statistic analysis. These differences may stem from methodological differences - the delta statistic detects a different type of evolutionary signal from RERconverge - but they may also be driven by potential bias in genes captured by the delta statistic as shown in Figure 9. In the case of the diet phenotype, the delta statistic was biased toward capturing signal driven by slower evolution of carnivore lineages compared to other lineages. Although the results are only explored for one trait, our results would suggest that the delta statistic may be biased toward detecting genes with patterns of evolution driven by single pairwise comparisons evolving in one direction. That limitation is confounded by a lack of post-hoc testing available through the delta statistic to evaluate which comparisons contribute to the omnibus signal.

Our methodological updates and analysis provides new insights into the evolution of diet. Throughout the course of the research, we created a ranked list of genes that have convergently evolved with diet across mammals (Data S1). These genes were enriched for pathways like “abnormal digestion” and “digestive tract development”, which suggest that there is substantial signal within the analysis (Data S2). Applying the categorical updates to RERconverge simultaneously maximizes the number of pathways detected and the proportion of diet-related pathways detected. However, additional analyses are required to create a more definitive reference. Future work may further improve our understanding of dietary evolution by integrating larger sets of species and more fine-scale resolution of dietary phenotypes, beyond the categories we have included here.

As demonstrated by our comparisons of CTMM and previous binary ancestral reconstruction, the assignment of phenotypes to internal branches has a dramatic effect on results. As such, analyses on the specific effects of different reconstruction methods, particularly reconstruction with more granular diet phenotypes, would provide further insight into the evolution of the diets analyzed in this paper in addition to the evolution of the more granular phenotypes. Additionally, diet can be represented in ways other than a non-ordinal categorical trait, such as the varying degrees of carnivory expressed by different mammals (Wilman et al. 2014; Pollard, Meyer, and Puckett 2023). Evaluation of diet from the continuous perspective and comparing those results with categorical analyses would provide confidence and nuance to the genes and pathways connected with diet evolution. Further, to confidently connect the associations with the evolution of diet, examination with other tools (Hyphy, BUSTED-ph) should be performed,and functional validation of genes detected through this analysis would be ideal.

There are already many applications of tests for convergent evolution across a variety of genomes (Marcovitz et al. 2019). The reduction in cost of genome sequencing has led to an encouraging proliferation of comparative genomics resources. In recent years, these have expanded from dozens of species (Zhang et al. 2014; Hecker and Hiller 2020) to hundreds of species (Christmas et al. 2023). Additional coordinated large scale genome sequencing efforts continue to expand resources and genome quality (Rhie et al. 2021; Teeling et al. 2018). With these resources, new computational approaches are being developed to obtain more accurate protein annotations from these diverse genomes (Kirilenko et al. 2023). In parallel, field work and coordinated databases are providing increased high-quality data on trait annotations. As the genome sequences, protein annotation, and trait annotations continue to improve, there will be an increasing need for methods that can handle increased scale of data and trait complexity. The improved ability of RERconverge to handle categorical traits provides a good foundation for the community to use newly emerging resources to make new biological discoveries.

## Methods

### Mammalian Coding Sequence Alignments and Trees

Mammalian coding sequence alignments were extracted from a whole-genome multiple alignment of 120 mammal species (Hecker and Hiller 2020). Coding exons were extracted based on human genome (hg38) coordinates from GENCODE version 36 (GENCODEV36) (Frankish et al. 2019). Transcripts were chosen only from “protein_coding” genes and the “canonical” isoforms were used, that is a single isoform meant to represent each gene. Exon alignments were extracted from a maf format alignment with human as reference using functions from the RPHAST package (Hubisz, Pollard, and Siepel 2011) . RPHAST was also used to enforce the human reading frame and translate all sequences to amino acids. The resulting amino acid alignments were used to infer gene trees for each gene using a fixed species tree topology using the phylogenetic package phangorn (Schliep 2011).

### Diet Phenotype Assignment

#### Phenotypes

Phenotypes used in the relaxation analysis were Herbivore (49 sp.), Omnivore (25 sp.), Insectivore (18 sp.), Carnivore (10 sp.), Piscivore (9 sp.), and Anthropivore (5 sp.); assigned according to methods described in the later section (Supplementary file 1 (phenotype vector); Supplementary file 2 (tree image)).

In addition, the grouping of Carnivore-Piscivore was combined to the phenotype Vertivore (19 sp.) and Omnivore-Anthropivore combined to the phenotype Generalist (30 sp.). Piscivores and anthropivores are subsets of carnivores and omnivores respectively, and thus provided biologically sound phenotype merges to allow for direct comparison between different numbers of categories used to describe the same species set.

Anthropivores are omnivorous species which can live in close association with humans, and derive food sources from non-typical human sources, such as cardboard or plaster. This does not necessarily reflect the species’ diet in the wild, but demonstrates a biologically-derived subset of omnivores. This subset of another phenotype was useful for direct comparisons in timing between category numbers, and was the purpose of this phenotype’s inclusion. As it was being used only in the capacity of being a subset of omnivores, anthropivory’s poor level of phenotype description was irrelevant.

#### Phenotype assignment

Diet data was collected from each species from Walker’s Book Of Mammals (Nowak 1999). Species which exclusively or nearly exclusively (90%+) consumed plant matter were considered Herbivores. Species which consumed exclusively or nearly exclusively arthropods were considered Insectivores. Species which consumed exclusively or nearly exclusively terrestrial vertebrates were considered Carnivores.

Species which exclusively consumed aquatic vertebrates, shellfish, and/or bivalves were considered Piscivores. Further distinction between aquatic vertebrates and invertebrates (mirroring the carnivore-insectivore split) was considered, but not performed to maintain a reasonable category number and due to the small size of the potential aquatic-insectivore clade.

Species which consumed more than one type of food source were considered Omnivores. Species which were capable of surviving off of primarily human-derived food sources (eg. food waste, cardboard, plaster) were classified as Anthropivores. In the 6-way analysis, all 6 category types were considered. In the main three way analyses, the categories were compressed to only three groups in such a way to maintain all of the species in the dataset within the 3-way analysis. To this end, insectivores were considered carnivores since, not distinguishing between vertebrates and invertebrates, insectivores are a specialized carnivore. As the primary goal of the main 3-category analyses was to demonstrate the categorical method rather than to identify fine-grained genetic differences between diet types, this simplification was made. Additionally, this simplification affected fewer than 10 of the 115 species in the phylogeny. As a result, additionally, some species which were considered omnivores in the 6-category analysis were considered carnivores in the 3-category analyses, due to those species consuming a mixture of vertebrates and invertebrates, which are not distinguished in the 3-category analyses.

#### CTMC Ancestral Reconstruction

Categorical RERconverge uses a continuous time markov model of evolution in order to perform ancestral state reconstruction. This is an important step because it allows RERconverge to associate relative evolutionary rates with the convergent evolution of the phenotype across the entire phylogeny, not just extant branches. Maximum likelihood estimation is used to infer a transition rate matrix, Q, from the user supplied phylogeny, rate model, and extant phenotype data. The phylogeny is the master tree which includes all species in the analysis and has branch lengths representing average genome-wide evolutionary rates. The transition matrix is inferred using the *fit_mk* function from the castor package (Louca and Doebeli 2018), and is used to compute the marginal ancestral likelihoods at each node. Code for the computation of ancestral likelihoods is heavily based on *ace* from the package ape, using the same double pass algorithm but is modified to work with unrooted, non-dichotomous trees (Paradis, Claude, and Strimmer 2004). Each node is then assigned the state with the maximum marginal likelihood. Though this is not the same as the assignment of states that maximizes the joint likelihood, marginal ancestral likelihoods can be computed rapidly even for large phylogenies.

The rate model describes the number and position of free rate parameters in the transition rate matrix. In order to compare rate models, RERconverge implements a likelihood ratio test which computes the log likelihood of the fitted transition matrix under each user supplied rate model. Pairwise comparisons are made between each rate model with its more complex rate models and the likelihood ratio and p-value is computed for each comparison. If the two rate models are nested (the simpler one is a special case of the more complex one), then the likelihood ratio is distributed as a chi-squared with degrees of freedom equal to the difference in the number of free parameters between the simpler and more complex model and the p-value is determined accordingly (Pagel 1994). Other packages implement a similar test for nested models including *anova* in the ape package (Paradis et al., n.d.). However, the likelihood ratio test in RERconverge works with both nested and non-nested models and will automatically detect whether the models are nested or not. For non-nested models, Monte Carlo simulations are used to determine an empirical p-value for the likelihood ratio (Pagel 1994).

For binary phenotypes, the user can choose whether branches are considered foreground based on the state of possessing the convergent trait or based on the transition from not possessing to gaining the trait. Both methods were tested for hairless species with no large impact found (Kowalczyk, Chikina, and Clark 2022). However, with categorical traits especially when the ancestral trait is unclear, the number of possible transition types is quadratic with respect to the number of categories. Thus we decided to assign edges based on state rather than transition. Edges were assigned the state of their descendant nodes such that all edges leading to extant species are assigned based on the observed phenotype.

In order to handle missing species, RERconverge computes “paths” (Kowalczyk et al. 2019). When species are missing, certain nodes are no longer necessary so edges will be combined into “composite” edges and the paths describe what state values to use for these composite edges. Categorical RERconverge assigns the composite edges the state of the most recent edge in the master tree. This ensures that composite edges leading to extant species always use the observed state.

#### Permulations

Permulations is an important part of RERconverge because it allows users to calculate reliable p-values for the association of genomic elements with convergent traits despite unknown sources of dependence in the data leading to non-uniform null p-value distributions (Saputra et al. 2021). Permulations generate permuted phenotype trees that maintain the phylogenetic relationships in the data. For instance, binary permulations maintain the same number of foreground species and the same structural relationships between foreground species (Saputra et al. 2021).

Categorical permulations are accomplished slightly differently than binary and continuous permulations because the simulation step does not use the Brownian motion model. Instead, phenotypes are simulated from the continuous time markov model that was used to reconstruct the ancestral history of the trait. As with binary and continuous permulations, the simulation is based on a phylogeny with branch lengths representing the average genome-wide evolutionary rate along that branch. Next, three steps are taken to ensure that the permulated phenotype contains the same number of species with each trait value as the original phenotype:

1. Rejection sampling: any simulated phenotype in which there are not the same number of extant species with each trait value as the original phenotype is rejected.
2. Permutation of internal traits: the simulated values for internal species are ignored. Instead, the originally inferred internal trait values are permuted and assigned to internal species in the permulated phenotype. Assignment is weighted by the ancestral likelihoods calculated from the simulated tip values. This ensures that the initial permutation of internal traits is more optimal than a completely random shuffle.
3. Re-organize internal traits: a search technique similar to simulated annealing is used to reorganize the internal states relative to the simulated extant states to improve the likelihood of the permulated phenotype. This is accomplished through a series of swaps. Pairs of candidate nodes are suggested based on which internal nodes from step (2) disagree most with the ancestral likelihoods at that node. A swap is made, and if the swap improves the likelihood of the tree, then the swap is kept. To avoid getting stuck, the swap is also made with a small probability even if it does not improve the likelihood of the tree. This generates a plausible trait history that exactly matches trait category counts and has a comparable likelihood to the original simulation.

Algorithmically step 3 is computed as follows: Let *Q* be the instantaneous transition rate matrix that was fit on the phenotype data. Let *A* be the matrix of ancestral likelihoods where each row is a node and each column is a phenotype state. The initial likelihood of the tree after step 2 is computed.

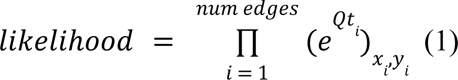

where *t*_*i*_ is the length of edge *i*, *e^Qt_i_^* is the transition probability matrix, *x*_*i*_ is the ancestor on edge *i*, and *y_i_* is the descendant on edge *i*. For each internal node _*i*_ currently assigned state *y*, ratios are computed for all other states *x*.

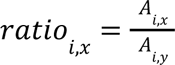

Thus a large ratio indicates that node *i* prefers to be in state *x* over its current state *y*.

A ratio, *r*1, is selected at random with more weight placed on larger ratios so that poorly assigned nodes are more likely to be swapped. Let *r*1 = *ratio_i,x_* where node *i* is currently in state *y*. Then, a second ratio, *r*2, is selected from a list of ratios, *ratio_j,y_*, where each node *j* is currently in state *x*. Once again, more weight is placed on larger ratios. A swap in which node _*i*_ is switched to state *x* and node *j* is switched to state *y* is then proposed.

The likelihood of the tree is recomputed under the swap. If the likelihood improves, then the swap is kept. If the likelihood does not improve, then the swap is made with probability 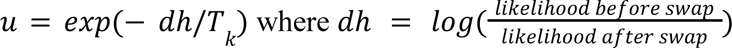. Thus if *dh* is large (the swap is very unfavorable), *u* will be small and it is less likely to make the swap. *T_k_* represents the “temperature” which is a common feature of simulated annealing algorithms. The temperature begins high during early iterations and decreases as the iterations go on. Thus, in the early iterations *u* is larger, and unfavorable are made with a higher probability. This allows the algorithm to try more swaps, especially early on, to increase the overall number of potential state configurations it explores, even if some of those swaps initially decrease the likelihood. Only allowing swaps that improve the likelihood may cause the trees to get stuck before they reach a more favorable state assignment.

At the end of each cycle *k*, the new temperature is calculated as 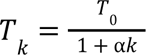 where *T*_0_ and α were chosen semi-arbitrarily and are currently 10 and 0.9 respectively. 100 cycles were run, each with 10 iterations for a total of 1000 iterations.

In order to determine the effectiveness and importance of each step in the above algorithm, we plotted the distributions of tree log likelihoods after each step (**Fig. S4**). After step 2, the log likelihoods decrease because we replace the simulated internal states with a permutation of the original internal states. However, after step 3, the completed permulated trees (tan) return to having the same or improved likelihoods compared to the simulations (blue). Thus, plotting the log likelihood distributions after each step demonstrates why step 3 is extremely important for ensuring the overall plausibility of the permulated phenotypes and thus maintenance of the types of phylogenetic dependencies present in the original phenotype tree.

Two different methods were originally developed for permulations; the CC (complete case) method which generates permulated phenotypes of the complete topology and SSM (species subset match) which is more time intensive in order to correct for missing species among some gene trees (Saputra et al. 2021). Since CC is significantly faster, we only implemented a categorical approach for the CC method.

Once the permulated phenotypes are generated, the correlation between RERs and each permulated phenotype is computed for each gene. Empirical p-values are then computed as the number of null effect sizes more extreme than the observed effect size for that gene. One sided empirical p-values are computed for the omnibus test and two-sided empirical p-values are computed for the pairwise tests.

### Permulations Timing

#### Timing permulation parameters

Permulations were performed using the same species as the main analysis, using all (19137) of the gene trees in the dataset. Permulations timing tests were performed with rp =”auto”, and ntrees = 5. This means that each time value is the time required to run 5 permulations, to account for fluctuations in each permulation’s generation time. Permulations were performed using both the SYM and ER rate models, for which the timing patterns broadly matched. The ARD rate model was not used, as it was not compatible with all of the phenotype sets. Trends were consistent across rate models, with the ER method being moderately faster than the SYM method, with the difference in speed between ER and SYM varying slightly with relaxation, category set, and category number. Permulations were performed on all combinations of phenotypes, including the merged Vertivore and Generalist phenotypes, for a total of 24 two-category comparisons, 34 three-category comparisons, 24 four-category comparisons, 8 five-category comparisons, and 1 six-category comparison, for a grand total of 91 category sets. For each category set, timing tests were performed five times, for a total of 25 permulations. All timing measurements presented outside the supplement were performed using the SYM model at 10% relaxation, as performing 5 or 6 category comparisons at 5% and 0% relaxation was non-practical (72+ hours) even when performing only 25 permulations. Data from the ER model follows similar trends, and the results for the ER model can be found in the supplement. Outliers were classified as any phenotype for which the time’s average z-score (inclusive of outlier) > 2, while the dataset’s standard deviation was greater than its mean. This allowed for detection of phenotypes which were dramatically affecting the mean of the dataset. All code used for permulation timing can be found here: [https://github.com/MichaelTene7/CategoricalPermulationsTiming]

#### Species number effect on permulations times

Tests for the effects of the number of species on permulation time were conducted by comparing the number of species in the tree to the permulation time for each number of categories. To account for differences caused by an increased number of categories, trees were only compared to others with the same number of categories.

For each number of categories, the effect of species number on permulation time was computed using linear regression. Calculations were performed both including and excluding outliers believed to be caused by phenotype clustering effects. Phenotype cluster effects are a phenomena observed where permulations take a much longer time if a single phenotype which comprises a large portion of the tree occurs in a monophyletic or nearly monophyletic clade; this results in very few valid alternates for use as a permulated tree, greatly increasing run time.

#### Category number effect

Category number effect on permulation time was determined by comparing the time of permulations with the categories split against the time of permulations where the categories were merged. For instance, the time for the four-way comparison Herbivore-Insectivore-Carnivore-Piscivore was compared against the three-way Herbivore-Insectivore-<Carnivore&Piscivore>. This comparison was done for all possible combinations of the merged categories, with 3vs2 categories (10 comparisons), 4vs3 categories (16 comparisons), 5vs4 categories (10 comparisons), and 6 vs 5 categories (2 comparisons). Additionally, comparisons were made where both of the mergeable categories were combined. For instance, the five-way Herbivore-Carnivore-Piscivore-Omnivore-Anthropivore was compared against the three-way Herbivore--. This was done for all 4vs2 categories (1 comparison), 5vs3 categories (2 comparisons), and 6vs4 categories (1 comparison). Finally, the overall averages of phenotype sets with each category number were compared to the phenotype set averages of the sets with each other category number.

#### Permulations Relaxation

Increasing the number of categories beyond three was demonstrated to significantly slow down permulations. The computational bottleneck occurs in the first step when phenotypes are simulated from the CTMM and simulations without the same number of extant species in each category as the original phenotype are rejected. Understandably, exactly matching counts becomes more challenging and thus slower as the number of categories increases. In order to handle these cases, a relaxed version of permulations was developed. This relaxation defines a percentage of the original counts that the simulated phenotype can fall within, and only rejects the simulation if the counts are not within this range. There are certain theoretical limitations to the current relaxation approach in that the internal category counts are still matched strictly to the original data. However, in practice we found that using a relatively small relaxation (of 10%) led to a significant speed improvement without reducing the evolutionary plausibility of the permulated phenotypes (as measured by the tree likelihoods) or causing large deviations in the p-values.

#### Relaxation effect on permulation speed

The full set of permulations were performed at relaxation levels of 0%, 5%, 10%, and 20% for the ER and SYM rate models, for all 91 category sets. Results presented are drawn from the SYM model, with the similar ER model results included in the supplement. For the 0% and 5% relaxation level on the 5 category and 6 category data, 10 permulations of each category set were unable to be completed within 72 hours. As such, the mean time is displayed as a conservative minimum by assuming the set of permulations took 72 hours to run to completion, though the real number is likely far higher, as in fact only approximately half the permulations completed in that time.

We compared the time to complete each of the 91 category sets between the relaxation levels for each rate model. Additionally, we compared the time to complete each of number-of-category permulations, eg. time to complete all three-way category sets, between relaxation levels.

#### Relaxation effects on permulation results

Permulation P-value calculations were performed on both unrelaxed and 10% relaxed category sets, with 20,000 permulations of each. The resulting p-values for all 19137 genes in the Hiller dataset were then compared, and the mean difference between relaxed and unrelaxed p-values for all genes was calculated. The category sets used were the three-category sets: Herbivore-- category set (the largest set) and the Piscivore-Carnivore-Insectivore category set (the smallest set). Category sets with more than 3 categories were not considered, due to the impracticality of running unrelaxed permulations with more than 3 categories.

#### Categorical Statistical Analysis

Relative evolutionary rates are associated with the categorical phenotype in one of two ways. The default approach is to use a nonparametric Kruskal Wallis test followed by pairwise Dunn tests (Ogle et al. 2023). There is also an option to use an ANOVA test followed by pairwise Tukey tests (Foundation for Statistical Computing, n.d.). The Kruskal Wallis test reports the epsilon squared effect size computed as 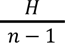 where *H* is the Kruskal Wallis test statistic and *n* is the total number of observations (species), and the corresponding p-value (King, Rosopa, and Minium 2018). The Dunn test reports the Z statistic for each pairwise test and the p-value after adjusting for multiple comparisons. The ANOVA test reports the eta squared effect size computed as 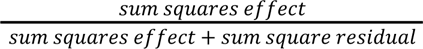, and the corresponding p-value. The Tukey test reports the difference between each pair of means and the p-value corrected for multiple comparisons.

### Other Methods

#### Binary RERconverge

We compared the results to those of multiple binary RERconverge analyses. We employed two approaches to account for there being greater than two categories. In the first approach, one category was chosen as the foreground and the other two categories were set as background. This was performed three times; once for each category. In the second approach, we ran six pairwise binary RERconverge analyses. In these analyses, only species from two categories were included, the other species were removed from the tree and were not included in the calculation of relative evolutionary rates. For each pair of categories, two analyses were performed by switching which category was considered foreground.

However, the second binary method with herbivore foreground was excluded from the analysis because it had over a 10x slower runtime than the other methods. The slowdown was caused by the rejection sampler because, to match foreground numbers, only permulated phenotypes including the same large clades as in the original data were being kept. This would incorrectly overestimate the inflation of low p-values.

Binary permulations with 10,000 permulated trees were performed on the remaining eight sets of binary gene correlations results. A modified version of binary permulations was performed which matches the exact extant and internal foreground species counts. Phylogenetic simulations are still used to replicate phylogenetic dependencies in the data but unlike original binary permulations, the permulated trees do not perfectly match the structure of the original phenotype due to greater complexity of the diet phenotype as compared to the phenotypes for which binary permulations was originally developed.

Rank-based pathway enrichments were calculated using the Wilcoxon Rank Sum test as implemented in the RERconverge package. Enrichment permulations were performed by calculating null enrichment statistics on all 10,000 null sets of gene correlation statistics and computing empirical p-values for each pathway based on the number of null enrichment statistics as or more extreme than the observed enrichment statistic.

#### Phylogenetic Signal

The delta statistic of a phylogenetic signal measures the degree to which a trait evolves according to a given phylogeny. It is based on the idea that when a trait is highly associated with a phylogeny, that phylogeny will be good at predicting the ancestral states of species with low uncertainty because the phylogeny effectively traces the evolutionary history of the trait. Thus the delta statistic summarizes the amount of uncertainty, measured as Shannon entropy, in a given set of ancestral likelihoods (Borges et al. 2019).

We used the publicly available, faster, python code from a recent update to this paper (Ribeiro et al. 2023) to calculate the delta statistic for each of our 19,137 gene trees with the 3-category diet phenotype. Ancestral likelihoods were calculated using the *ace* function from the ape package with an all rates different (ARD) rate model in order to be consistent with Borges et al. 2019.

All of the gene trees used in the RERconverge analysis were confined to have the same topology as the master tree. We believe this constraint may limit the degree of variation in phylogenetic signal observed between different genes and impact the ability of the delta statistic to identify significant genes. Thus, we constructed unconstrained gene trees using iqtree (Nguyen et al. 2015) and repeated the delta statistic analysis.

Enrichments were calculated using the same approach as for all the RERconverge methods, using the Wilcoxon Rank Sum test as implemented in the RERconverge package. Enrichment permulations were not performed.

#### PhylANOVA

We used PhylANOVA from the phytools package to perform the phylogenetic simulation method as described by Garland et. al. 1993 (Garland et al. 1993; Revell 2012). This method is designed to correct p-values for non-independence between species due to hierarchical phylogenetic relationships. This is accomplished by computing p-values from empirical distributions of *F* statistics (the ANOVA statistic) rather than from standard tabular values. The empirical distributions of *F* statistics are determined from simulations of the continuously valued response variable (in this case the relative evolutionary rates) along a known phylogeny using a Brownian Motion model. The assignments of species to each category remains the same during the simulations and calculation of the null distribution of *F* statistics.

Technically, the relative evolutionary rates are not a continuously valued phenotype that evolved along the tree, rather they are a measure of evolutionary change calculated from the multiple sequence alignments used to construct the trees. However, the branch lengths we used for the simulations are those from the average tree rather than the individual gene trees from which the relative evolutionary rates are calculated so this should reduce the problem of circularity.

PhylANOVA was run with 10,000 simulations in order to be consistent with the number of permulations performed on the RERconverge methods.

Enrichments were calculated using the Wilcoxon Rank Sum test as implemented in the RERconverge package. Enrichment permulations were not performed.

## Supplementary Figures and Tables

**Figure S1.**
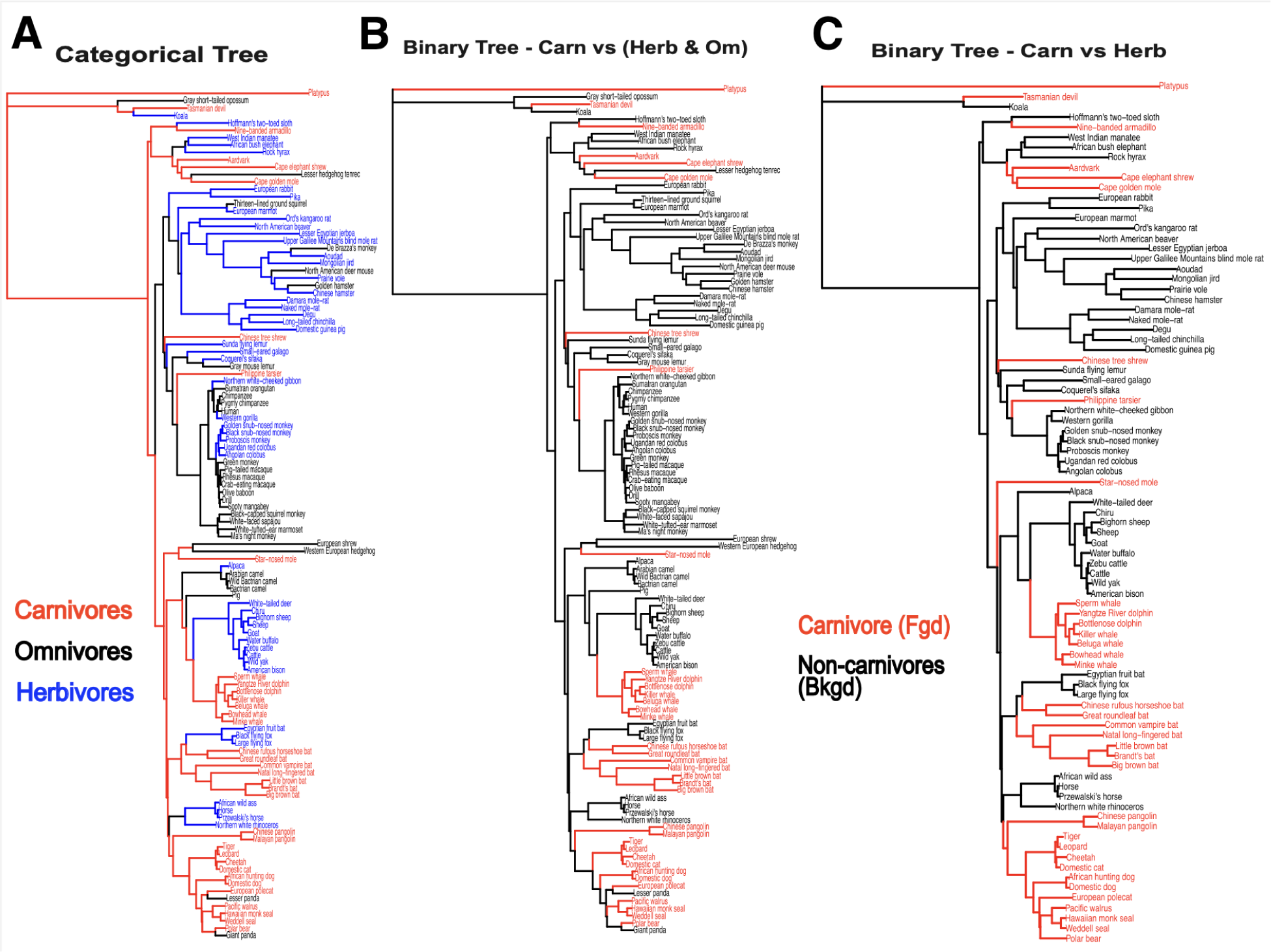
A) Categorical reconstruction using maximum likelihood applied to a continuous time markov model on the full phylogeny used in the analysis. B) Example of one of the binary reconstructions in which the foreground is carnivores and the background is herbivores and omnivores. Uses a maximum parsimony based approach. C) Example of one of the binary reconstructions in which the foreground is carnivore and the background is herbivores, with omnivores removed. Uses a maximum parsimony based approach.

**Figure S2.**
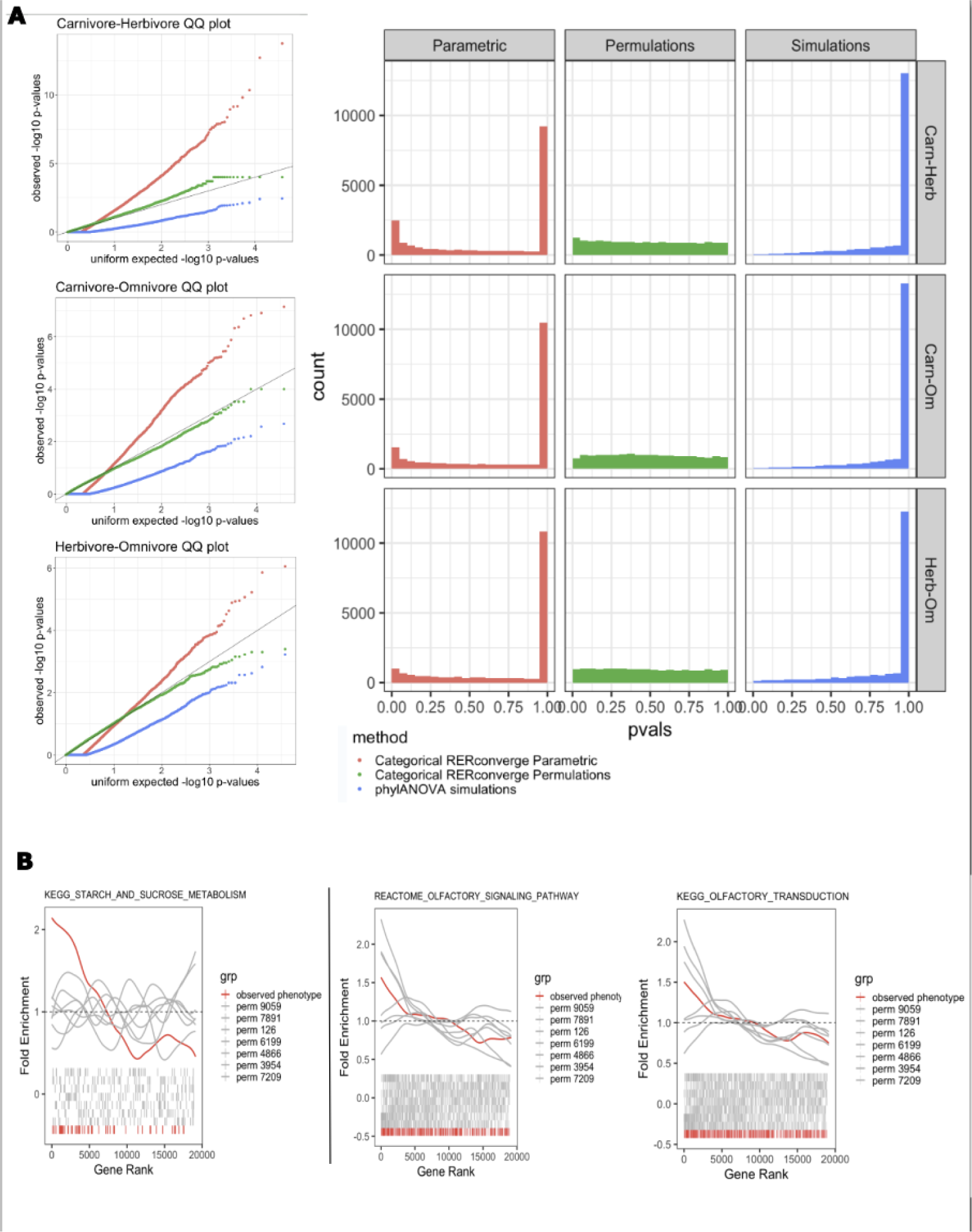
A) Quantile quantile plots and histograms of the, categorical RERconverge raw (parametric) p-values (red) and permulation p-values (green) and the phylanova simulation p-values (blue). B) Fold enrichment and barcode plots showing the enrichment of genes in the kegg starch and sucrose metabolism pathway (left) and two olfactory pathways (right). Red indicates the results for the observed phenotype, gray indicates the results for a random selection of seven (out of the 10,000) permulated phenotypes.

**Table S1.** Average time of all phenotype sets for each category number.

|  | Mean | SD | No Outlier Mean | No Outlier SD |
| --- | --- | --- | --- | --- |
| Two Category | 7.1987 | 0.428 | 7.198 | 0.428 |
| Three Category | 9.1044 | 0.951 | 9.1044 | 0.951 |
| Four Category | 38.591 | 34.49 | 32.614 | 18.64 |
| Five Category | 2358.0 | 5049. | 302.38 | 214.1 |
| Six Category | 4933.0 | NA | 4933.0 | NA |

**Table S2.** Percent speed increases for reducing the number of categories in the analysis, at each relaxation level.

| Relaxation (%) | comparison | Mean Difference (%) | Sd Difference (%) | Outlier Mean Difference (%) | Outlier Sd Difference (%) |
| --- | --- | --- | --- | --- | --- |
| Single | Single | Single | Single | Single | Single |
| 20 | 3v2 | 2.510765 | 3.333216 | 2.510765 | 3.333216 |
| 20 | 4v3 | 32.56813 | 33.64818 | 32.56813 | 33.64818 |
| 20 | 5v4 | 155.0417 | 196.6984 | 355.9884 | 661.9572 |
| 20 | 6v5 | 320.1345 | 385.5756 | 320.1345 | 385.5756 |
| 10 | 3v2 | 23.12227 | 13.83585 | 23.12227 | 13.83585 |
| 10 | 4v3 | 284.4025 | 144.2443 | 376.9865 | 395.6866 |
| 10 | 5v4 | 1501.997 | 762.154 | 4703.116 | 6873.465 |
| 10 | 6v5 | 1715.987 | 2268.198 | 1715.987 | 2268.198 |
| 5 | 3v2 | 48.57902 | 27.11924 | 48.57902 | 27.11924 |
| 5 | 4v3 | 1195.116 | 835.5254 | 1195.116 | 835.5254 |
| 5 | 5v4 | NaN | NA | NaN | NA |
| 5 | 6v5 | NaN | NA | NaN | NA |
| 0 | 3v2 | 51.14266 | 30.8599 | 51.14266 | 30.8599 |
| 0 | 4v3 | 1591.495 | 1627.517 | 1591.495 | 1627.517 |
| 0 | 5v4 | NaN | NA | NaN | NA |
| 0 | 6v5 | NaN | NA | NaN | NA |
| Double | Double | Double | Double | Double | Double |
| 20 | 4v2 | 32.80652 | NA | 32.80652 | NA |
| 20 | 5v3 | 163.9363 | 88.87492 | 163.9363 | 88.87492 |
| 20 | 6v4 | 1028.2 | NA | 1028.2 | NA |
| 10 | 4v2 | 481.8241 | NA | 481.8241 | NA |
| 10 | 5v3 | 5014.968 | 3484.625 | 5014.968 | 3484.625 |
| 10 | 6v4 | 32220.14 | NA | 32220.14 | NA |
| 5 | 4v2 | 1947.827 | NA | 1947.827 | NA |
| 5 | 5v3 | NaN | NA | NaN | NA |
| 5 | 6v4 | NaN | NA | NaN | NA |
| 0 | 4v2 | 1173.703 | NA | 1173.703 | NA |
| 0 | 5v3 | NaN | NA | NaN | NA |
| 0 | 6v4 | NaN | NA | NaN | NA |

**Table S3.** Table of the time average and standard deviations of time required to complete a phenotype set with a given number of categories, at each relaxation level tested. %Dif column represents the percentage speed increase of the relaxation level over 0% relaxation. As times for 0% and 5% relaxation for could not be completed, times were very conservatively estimated as the minimum time required to reach the run time produced (see methods), and no %Dif could be created.

|  | Mean_0 | Sd_0 | Mean_5 | Sd_5 | %Dif_5 | Mean_10 | Sd_10 | %Dif_10 | Mean_20 | Sd_20 | %Dif_20 |
| --- | --- | --- | --- | --- | --- | --- | --- | --- | --- | --- | --- |
| twoCategory | 8.11 | 0.543 | 8.007 | 0.549 | 1.370 | 7.198 | 0.428 | 12.76 | 8.243 | 0.4890 | -1.53157 |
| threeCategory | 14.35389 | 4.1969 | 13.883 | 5.0644 | 3.3853 | 9.1044 | 0.951 | 57.657 | 8.8042 | 0.5050 | 63.033 |
| fourCategory | 243.393 | 112.52 | 205.41 | 162.61 | 18.486 | 32.614 | 18.64 | 646.26 | 11.6372 | 2.3934 | 1991.49 |
| fiveCategory | 4320+ | NA | 4320+ | NA | NA | 302.38 | 214.1 | NA | 28.408 | 23.735 | NA |
| sixCategory | 4320+ | NA | 4320+ | NA | NA | 4933.0 | NA | NA | 119.679 | NA | NA |
| twoCategoryAll | 8.11748 | 0.543 | 8.0077 | 0.5498 | 1.3704 | 7.1987 | 0.428 | 12.762 | 8.2437 | 0.4890 | -1.5315 |
| threeCategoryAll | 14.3538 | 4.1969 | 13.883 | 5.0644 | 3.3853 | 9.1044 | 0.951 | 57.657 | 8.8042 | 0.5050 | 63.033 |
| fourCategoryAll | 276.698 | 196.8 | 205.41 | 162.61 | 34.70 | 38.591 | 34.49 | 616.98 | 11.6372 | 2.3934 | 2277.6 |
| fiveCategoryAll | 4320+ | NA | 4320+ | NA | NA | 2358.0 | 5049. | NA | 61.5213 | 96.201 | NA |
| sixCategoryAll | 4320+ | NA | 4320+ | NA | NA | 4933.0 | NA | NA | 119.679 | NA | NA |

**Figure S3.**
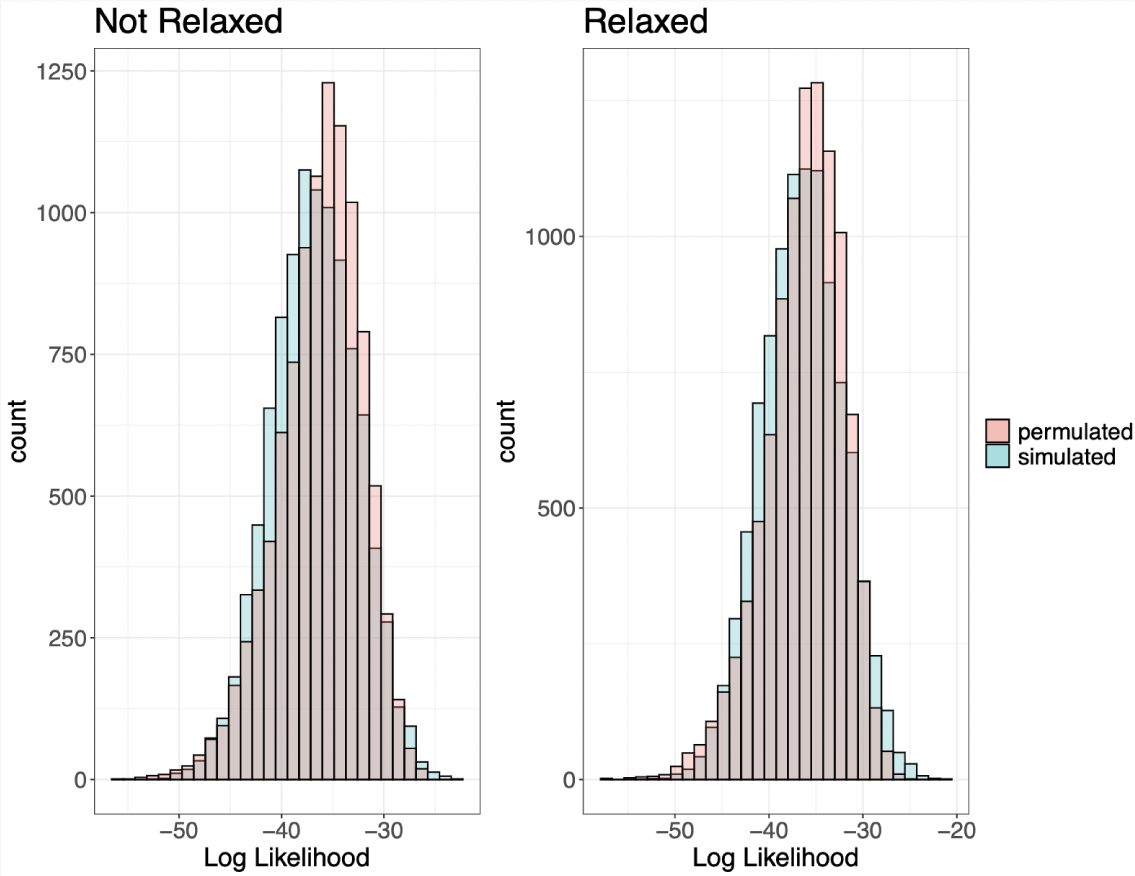
Histograms of the log likelihoods of the 10,000 simulated trees (blue) compared to the log likelihoods of the finished permulated trees (pink) for permulations without relaxation (left) and permulations with relaxation (right).

**Figure S4.**
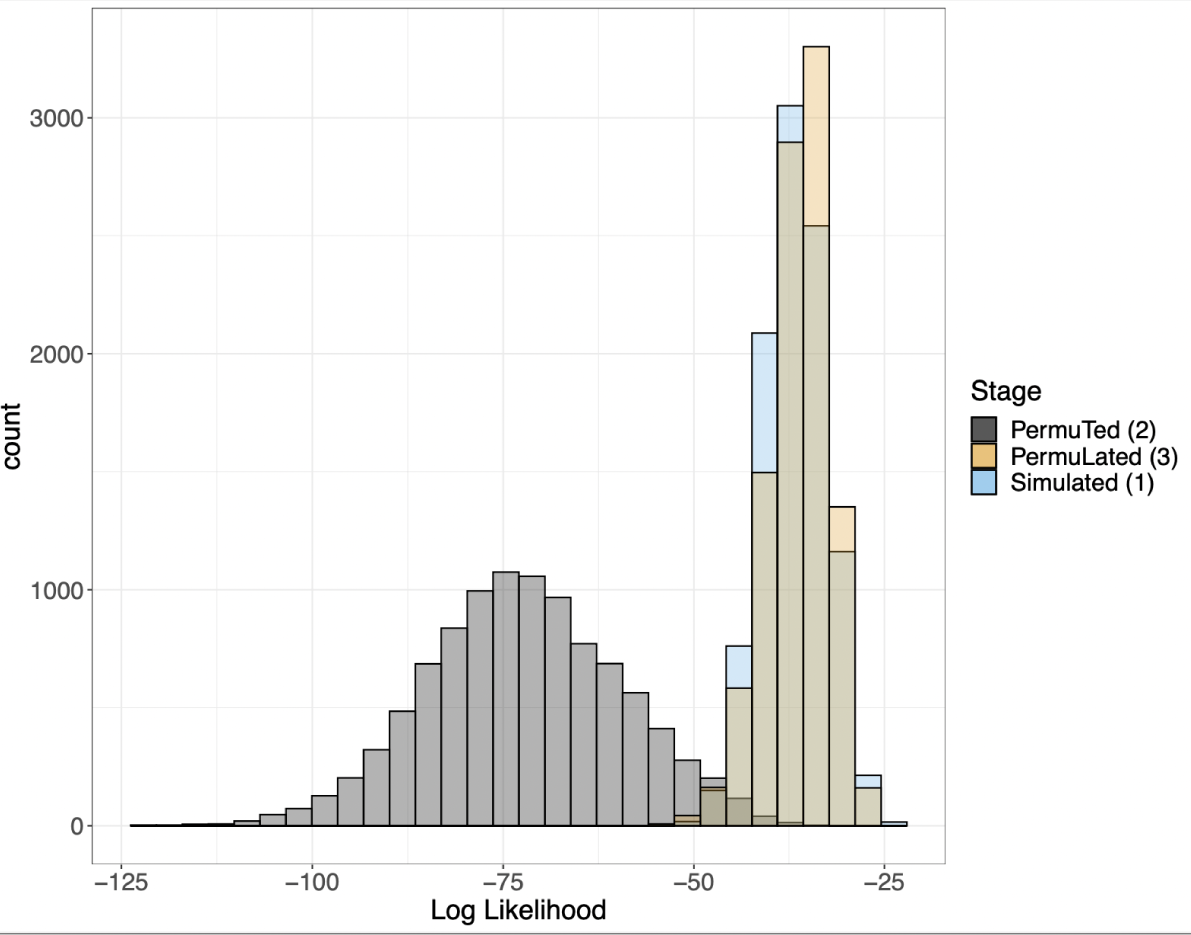
In blue are the log likelihoods of the 10,000 simulated trees (after step 1), in gray are the log likelihoods of the 10,000 permuted trees (after step 2), and in tan are the log likelihoods of the 10,000 finished permulated trees (after step 3).

**Fig S5.**
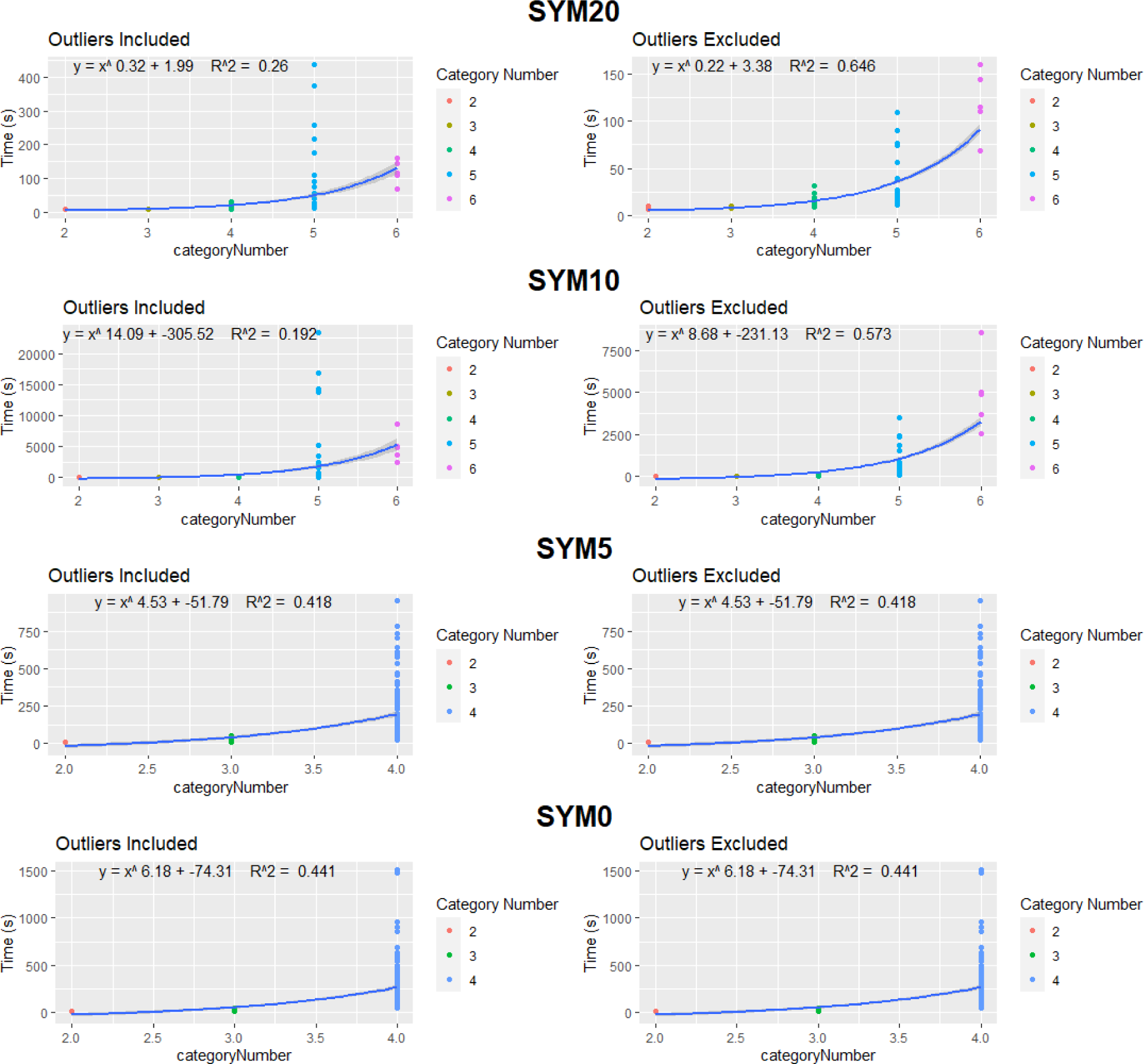
Graphs of the exponential effect of category number on permulations time. Note that the 5% and 0% relaxation graphs do not include 5 category or 6 category analyses.

## Acknowledgements

Portions of this research were conducted on Lehigh University’s Research Computing infrastructure partially supported by NSF Award 2019035. This material is based upon work supported by the National Science Foundation under Grant No. 2233124 (to WKM).

This research was supported in part by the University of Pittsburgh Center for Research Computing, RRID:SCR_022735, through the resources provided. Specifically, this work used the HTC cluster, which is supported by NIH award number S10OD028483.

Summer Undergraduate Research Fellowship (SURF)

Carnegie Mellon Neuroscience Institute Postdoctoral Fellowship

NSF 2046550

2R01 HG009299-06A1

NSF 2238125

