## Supplementary figures and images for "RERconverge Expansion: Using Relative Evolutionary Rates to Study Complex Categorical Trait Evolution"

### Supplemental file 2

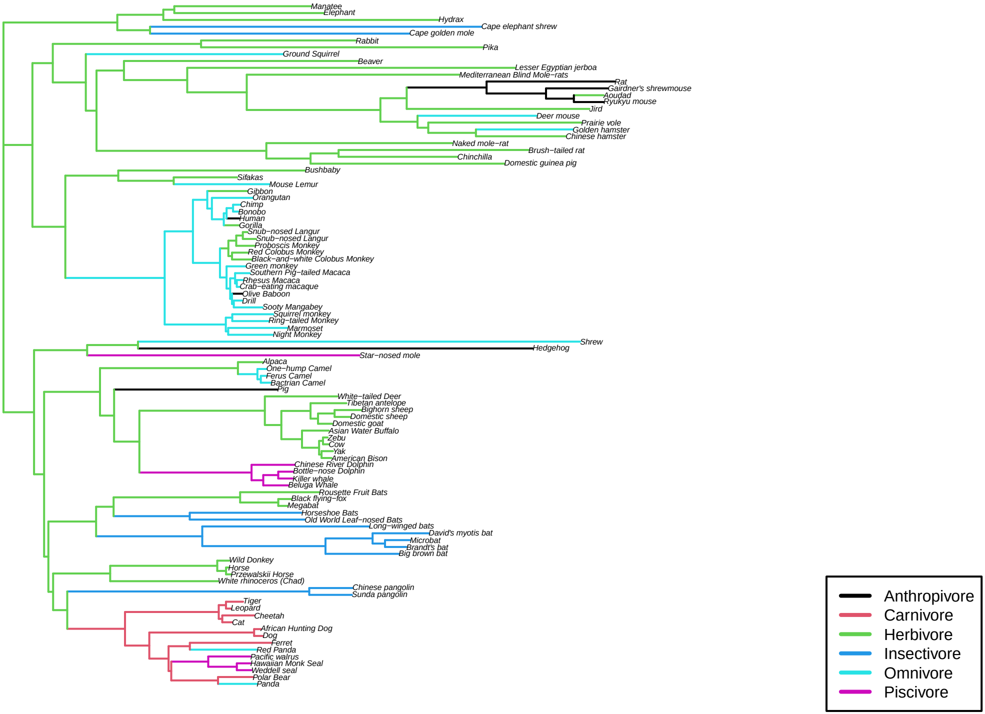

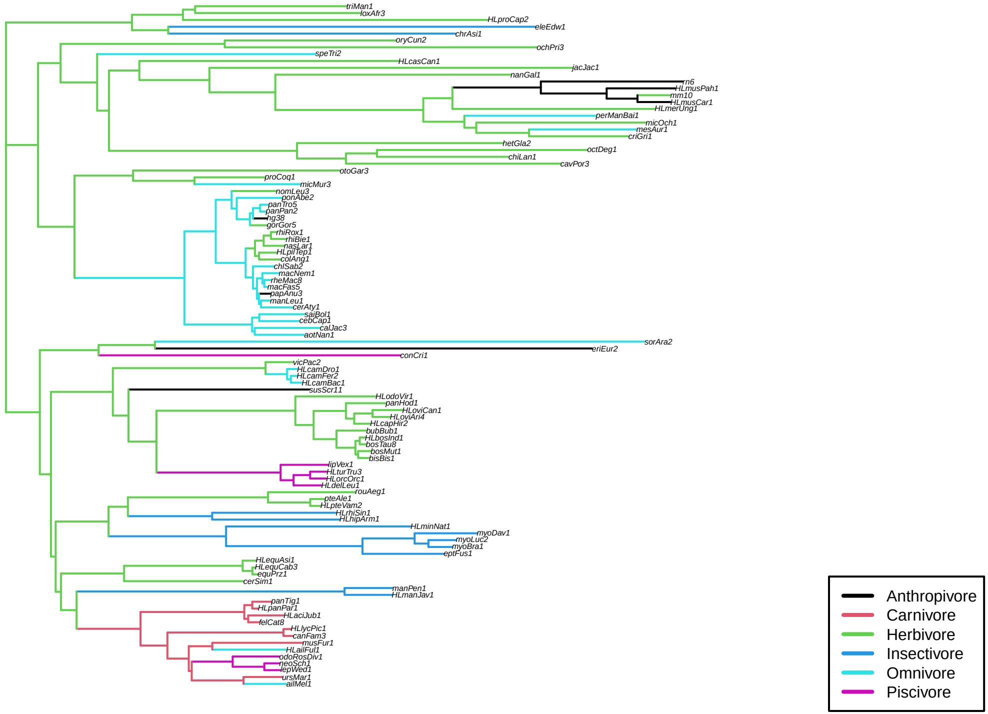
